# Integration of Brain and Behavior Measures for Identification of Data-Driven Groups Cutting Across Children with ASD, ADHD, or OCD

**DOI:** 10.1101/2020.02.11.944744

**Authors:** Grace R. Jacobs, Aristotle N. Voineskos, Colin Hawco, Laura Stefanik, Natalie J. Forde, Erin W. Dickie, Meng-Chuan Lai, Peter Szatmari, Russell Schachar, Jennifer Crosbie, Paul D. Arnold, Anna Goldenberg, Lauren Erdman, Jason P. Lerch, Evdokia Anagnostou, Stephanie H. Ameis

**Author notes:** **Corresponding Author:** Stephanie H. Ameis, Centre for Addiction and Mental Health 80 Workman Way, 5224, Toronto, ON, Canada, M6J 1H4.

## Abstract

Autism spectrum disorder (ASD), obsessive-compulsive disorder (OCD) and attention-deficit/hyperactivity disorder (ADHD) are clinically and biologically heterogeneous neurodevelopmental disorders (NDDs). The objective of the present study was to integrate brain imaging and behavioral measures to identify new brain-behavior subgroups cutting across these disorders. A subset of the data from the Province of Ontario Neurodevelopmental Disorder (POND) Network including participants with different NDDs (aged 6-16 years) that underwent cross-sectional T1-weighted and diffusion-weighted magnetic resonance imaging (MRI) scanning on the same 3T scanner, and behavioral/cognitive assessments was used. Similarity Network Fusion was applied to integrate cortical thickness, subcortical volume, white matter fractional anisotropy (FA), and behavioral measures in 176 children with ASD, ADHD or OCD with complete data that passed quality control. Normalized mutual information (NMI) was used to determine top contributing model features. Bootstrapping, out-of-model outcome measures and supervised machine learning were each used to examine stability and evaluate the new groups. Cortical thickness in socio-emotional and attention/executive networks and inattention symptoms comprised the top ten features driving participant similarity and differences between four transdiagnostic groups. Subcortical volumes (pallidum, nucleus accumbens, thalamus) were also different among groups, although white matter FA showed limited differences. Features driving participant similarity remained stable across resampling, and the new groups showed significantly different scores on everyday adaptive functioning. Our findings open the possibility of studying new data-driven groups that represent children with NDDs more similar to each other than others within their own diagnostic group. Such new groups can be evaluated longitudinally for prognostic utility and could be stratified for clinical trials targeted toward each group’s unique brain and behavioral profiles.

## INTRODUCTION

Neurodevelopmental disorders (NDDs), such as autism spectrum disorder (ASD), pediatric obsessive-compulsive disorder (OCD) and attention-deficit/hyperactivity disorder (ADHD), are often associated with poor cognitive and functional outcomes, although long-term trajectories vary considerably[1–4]. There are high rates of co-occurrence between different NDDs[5], as well as similarities in functional impairment[6] and clinical features (e.g. inattention[7], repetitive behaviours[5]). Together with similarity in genetic variants implicated in risk across NDDs[8], these convergences suggest that some children with different NDD diagnoses may be more similar to each other at the biological and behavioral level despite current distinct categorical (i.e., DSM-5/ICD-10-based) classifications.

Recent transdiagnostic neuroimaging studies highlight the heterogeneity within and across different NDDs and emphasize the need for new research models to move the field forward[9–11]. For example, a prior study from our group using the transdiagnostic Province of Ontario Neurodevelopmental Disorders (POND) dataset showed that children with ASD, ADHD, or OCD all featured non-distinct corpus callosum alterations compared to typically developing controls[12]. A continuous positive association between white matter microstructure and adaptive (everyday) functioning across children, irrespective of NDD category was also found. Others have also reported on the absence of clear biological distinctions on structural or functional neuroimaging measures when comparing different NDD diagnostic groups [13–18]

Data-driven clustering approaches offer a methodological alternative to conventional comparisons between clinically defined groups. This alternative approach may better disentangle heterogeneity within and across current diagnostic categories to identify participant subgroups that may be more similar to each other in brain or behavior[10]. Some of these approaches use data integration techniques to identify data-driven subgroups beyond using neuroimaging[19] or behavioral features alone [13]. Different clustering techniques can identify subgroups of participants across disorders with more similar brain-behavior profiles than those within a disorder[20,21], including a recent effort in the POND sample that integrated cortical thickness and behavioral measures, showing that identified clusters did not divide along diagnostic boundaries[22].

The present study aims to build on these efforts by simultaneously integrating different brain imaging phenotypes (regional cortical thickness, subcortical volume, and white matter tract fractional anisotropy, FA) with behavioral measures in children with primary ASD, ADHD or OCD clinical diagnoses using Similarity Network Fusion (SNF), a data integration approach[23]. SNF identifies participant similarity networks by integrating within and across data types, and thus groups participants together who are most similar to each other. We hypothesized that we would find new groups, each comprised of children with different NDDs who would show similar brain imaging and behavioral features to each other (i.e. within group) but different from other participants (i.e. between group); further, these differences would be of larger effect size than those found using categorical NDD diagnoses. We then examined whether differences between new groups would extend to out-of-model measures (e.g. functioning), hypothesizing again that a similar pattern would emerge. Finally, we examined the stability of our model, and explored whether supervised machine learning could be used to compare accuracy of subgroup identification using different sets of top contributing model features.

## MATERIALS AND METHODS

### Participants

Participants included children recruited through the POND Network between June 2012 to July 2017 from the Hospital for Sick Children and Holland Bloorview Kids Rehabilitation Hospital who were all scanned on the same Siemens Tim Trio (Malvern, Pa.) 3T magnetic resonance imaging (MRI) scanner located at the Hospital for Sick Children (Toronto, Canada). Additional data collection through POND is ongoing post scanner upgrade to the PrismaFIT, which was not analyzed for this report. Each institution received approval for this study from their respective research ethics boards. Following a complete description of the study, written informed consent/assent from primary caregivers/participants was obtained. Inclusion criteria included: age<18 years, presence of a primary clinical diagnosis of ASD, ADHD or OCD, confirmed using the Autism Diagnostic Interview-Revised[24] and Autism Diagnostic Observation Schedule-2[25] for ASD, Parent Interview for Child Symptoms[26] for ADHD, and the Schedule for Affective Disorders–Children’s Version (Kiddie-SADS)[27] and the Children’s Yale-Brown Obsessive Compulsive Scale[28] for OCD. Full-scale IQ was estimated using age-appropriate Wechsler scales in all participants. After quality control of MRI data (n=57 removed), removal of participants with missing behavioral data (n=26) and those older than 16 years (n=7) to ensure similar age variance across groups (Figure S1), data from a total of 176 participants were used for the main analyses (Table 1; see Supplementary Materials and Methods).

**Table 1.**
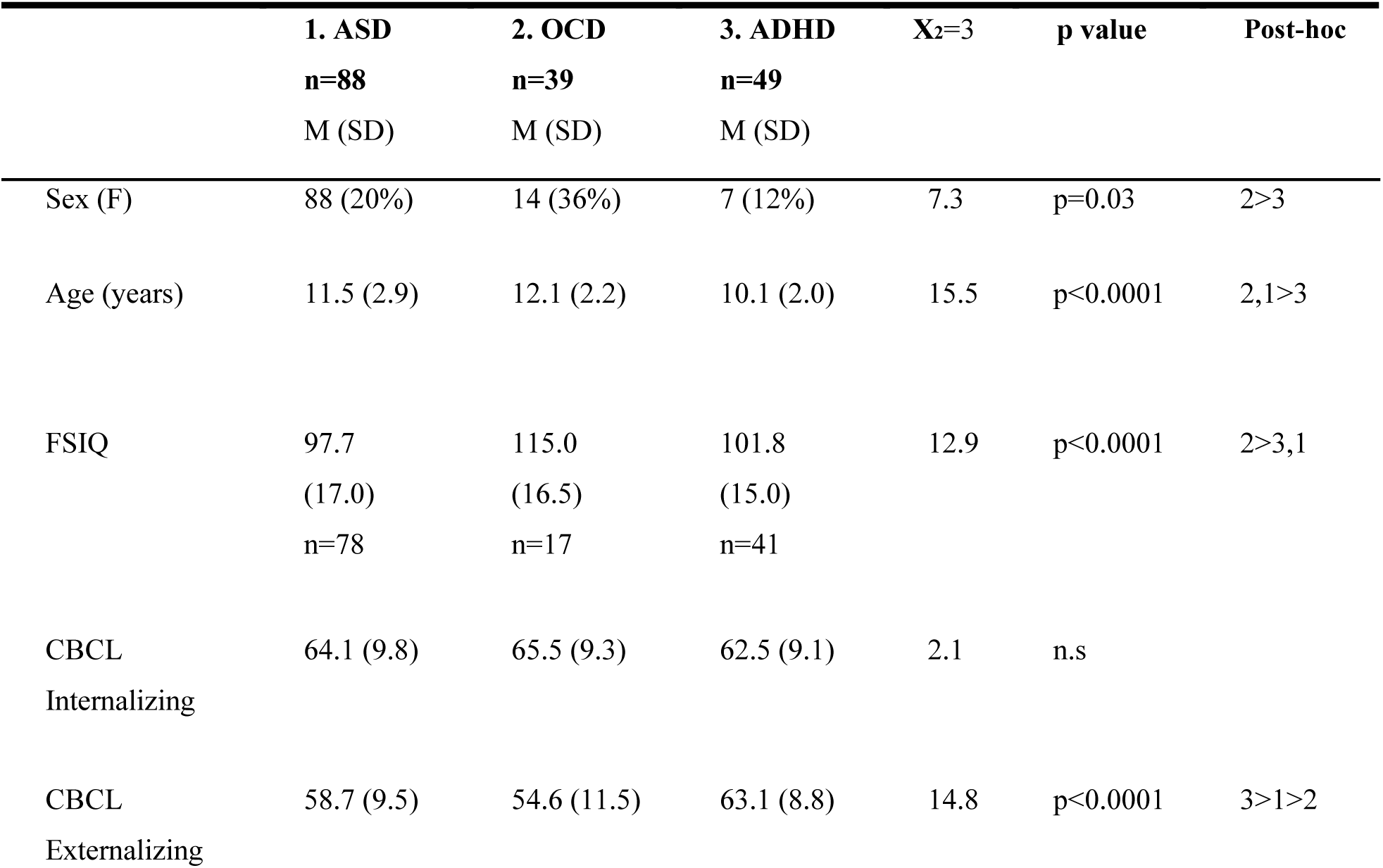
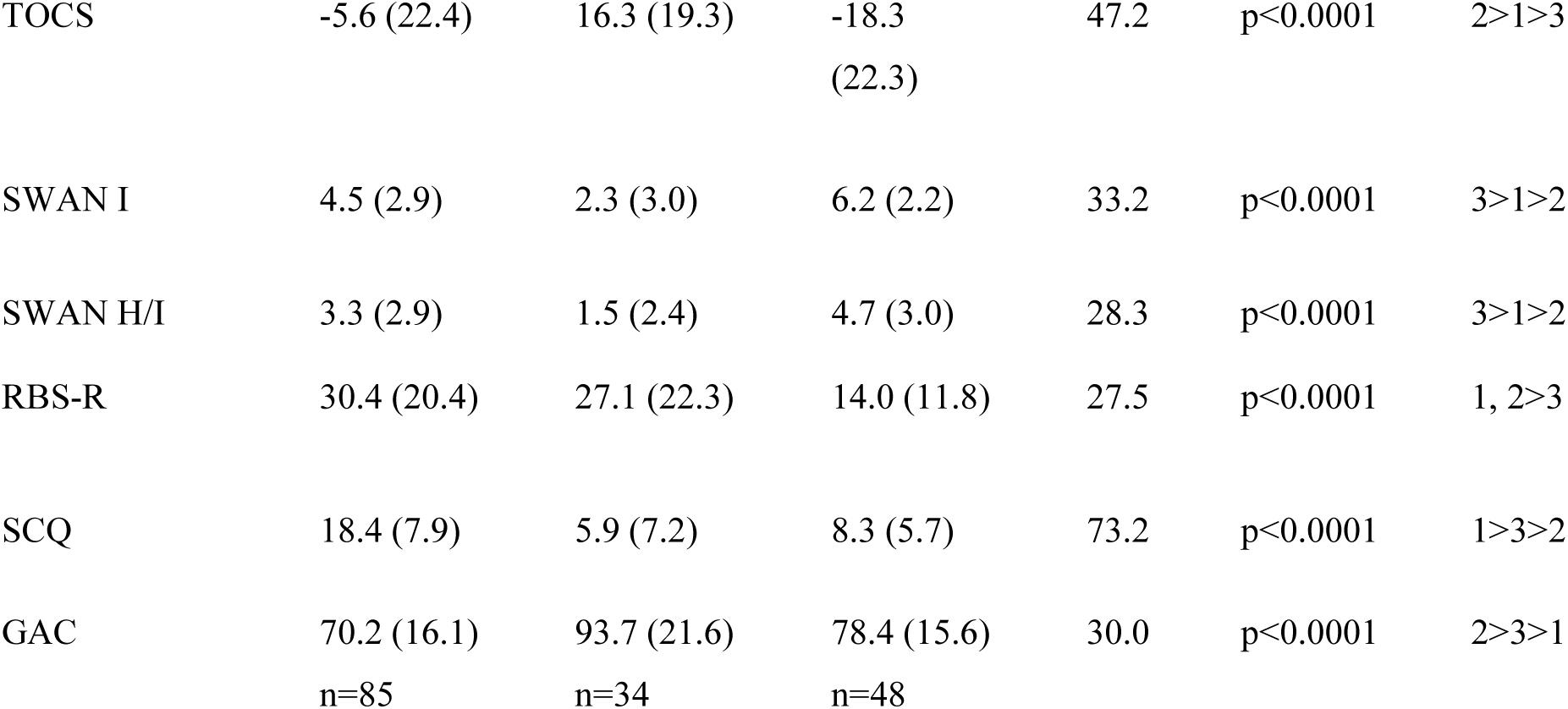
Sample characteristics presented by diagnostic group

### Clinical/Behavioral Assessments

Behavioral measures were selected to capture clinical features of each NDD that varied dimensionally across participants. Seven total raw parent-report scores from five behavioral scales were selected as input features for SNF analysis: the Child Behavior Checklist (CBCL 6-18) externalizing and internalizing broad-band scores[29]; Toronto Obsessive-Compulsive Scale (TOCS) total score[30]; Social Communication Questionnaire (SCQ) total score[31]; Repetitive Behaviors Scale-Revised(RBS-R) total score[32]; and total Strengths and Weaknesses of ADHD Symptoms and Normal Behaviour (SWAN) inattention and hyperactivity/impulsivity item scores[33]. The parent-reported general adaptive composite score from the Adaptive Behaviour Assessment System-II(ABAS-II)[34] capturing cross-disorder impairments in adaptive functioning was also assessed in participants.

All descriptive statistics are represented as means with standard deviations indicated in brackets except for self-reported sex, where percentage of females is included in brackets. Chi-squared (sex) and Kruskal-Wallis tests (all other measures) were used to test group differences. All participants were included in statistics except in the case of IQ and ABAS-II where a subset of the sample was used with available data (as indicated). ASD=Autism Spectrum Disorder, OCD=Obsessive Compulsive Disorder, ADHD=Attention-Deficit/Hyperactivity Disorder, F=female, FSIQ=Full-Scale Intellectual Quotient (estimated using WASI=Wechsler Abbreviated Scale of Intelligence or WISC=Wechsler Intelligence Scale for Children), CBCL Internalizing/Externalizing=Child Behaviour Checklist Internalizing or Externalizing t-score. Raw scores for Toronto Obsessive-Compulsive Scale (TOCS), Strengths and Weaknesses of Attention-Deficit/Hyperactivity-symptoms and Normal Behaviours (SWAN) inattention (I) and hyperactivity/impulsivity (H/I), Repetitive Behaviours-Revised (RBS-R), and Social Communication Questionnaire (SCQ) presented. The General Adaptive Composite (GAC) Scaled Score for the Adaptive Behaviour Assessment System-II (ABAS-II) is also presented.

### MRI Acquisition Parameters

All MRI data was acquired on the same 3T scanner using a 12-channel head coil. T1-weighted and diffusion-weighted acquisitions are detailed in Supplementary Materials and Methods [12,17,35].

### Image Analysis

Cortical thickness and surface area values (for 68 regions) and subcortical volumes (for 14 regions) were derived from T1-weighted images using the Desikan-Killiany Atlas in FreeSurfer (v.5.3)[36] for regional parcellation. Diffusion MRI data were preprocessed using a combination of FSL(https://fsl.fmrib.ox.ac.uk/fsl/fslwiki/) and MRtrix(http://www.mrtrix.org/) to denoise, upsample the data, and correct for motion and eddy currents (Supplementary Materials and Methods). Tract-based spatial statistics(v1.2)[37] and the ENIGMA DTI pipeline were used to estimate white matter FA as an indirect index of white matter microstructure (for 46 regions) [38]. Imaging metrics for subcortical volume, cortical thickness and white matter FA were included as brain-based input features in the SNF analysis. Subcortical volumes were divided by intracranial volume. Surface area was used subsequently as an out-of-model measure to test data-driven group differences

### Quality Control

Please see Supplementary Materials and Methods for quantitative and qualitative MRI quality control details.

### SNF Data Integration and Cluster Determination

SNF(v2.3.0 R package) was used to integrate structural imaging (cortical thickness, subcortical volume, white matter FA) and behavioral (CBCL, SCQ, RBS-R, SWAN, TOCS scores) data types (see Table S3 listing all SNF input features). Separate networks describing participant similarity for each data type were first created, followed by the use of a nonlinear combination method to iteratively fuse networks for each data type into a single participant similarity network representing the full spectrum of included features[23]. Similarity matrices for each of the four data types (i.e., cortical thickness, subcortical volume, white matter FA and behavioral data) were calculated using Euclidean distance with a nearest neighbours value of 18 and normalization parameter of 0.8, based on consultation with developers and suggested nearest neighbour value of sample size/10. Normalized mutual information (NMI) was used as a metric describing the overlap in a similarity matrix created using any single model feature compared to the fused matrix created using all model features (NMI range 0-1). Features with higher NMI scores indicate greater contribution to participant similarity. Spectral clustering (SNF *spectralClustering* function) was then applied to delineate groups based on participant similarity matrices determined using 135 model features across 1000 iterations of resampling 80% of participants. A silhouette plot quantified the similarity between participants within a given group compared to participants in all other groups. The R package qgraph (v.1.6.1) was used for visualization of relative similarity among participants.

### Comparisons Among Identified Data-Driven Groups on Demographic, Cognitive and Top Contributing Model Features

Separate one-way ANCOVAs were conducted to examine whether data-driven groups differed on age, sex and IQ measures. Based on the results, all subsequent analyses used to evaluate data-driven group distinctions on model features covaried for age, sex and IQ. Separate one-way ANCOVAs were conducted to provide a standardized effect size estimate (using eta squared) of data-driven group distinctions for model features contributing to participant similarity, as well as for diagnostic groups. Correction for multiple comparisons was applied to all 135 features using a false discovery rate (FDR) of 5%. Where ANCOVAs were significant, follow-up Tukey comparison tests were run to determine distinctions between specified groupings.

### Evaluation of Clusters

#### Cluster Stability

Internal cluster reproducibility was evaluated via bootstrap resampling across 1000 iterations of 80% of participants to calculate stability measures. We used proportion of resampling (i.e. would top features consistently remain top features), the percentage of time that each participant clustered with each other participant, and an Adjusted Rand Index measure of the overlap between clusters to determine stability of the data-driven model.

#### Extension of Data-Driven Group Differences to Out-of-Model Features

Brain and behavioral measures that were excluded from SNF analysis (i.e. ABAS-II General Adaptive Composite, surface area) were compared using ANCOVAs to evaluate whether distinctions found between data-driven groups extended to out-of-model features. Similarly, cortical thickness brain network measures of global efficiency, network strength, and density were compared across a range of thresholds using permutation testing. All analyses accounted for the effects of age, sex, and IQ,

#### Comparison of Classification Accuracy Based on Different Selections of Top Contributing Features

A random forest machine learning algorithm was applied across 100 permutations of randomly resampled participants with an 80/20 train-test split to evaluate the reliability of group prediction using different sets of top contributing model features. Due to the limitations of testing classification within the sample used to identify initial groups (versus out-of-sample testing), this approach was considered exploratory to help understand the reliability of group identification and to determine which features might be more likely to accurately classify groups (see Supplementary Materials and Methods).

## RESULTS

### Top Ranking Features Contributing to Formation of Four Transdiagnostic Data-Driven Groups

The four transdiagnostic data-driven participant similarity groups identified using SNF and spectral clustering (Figure 1, Table S2) featured an average silhouette width of 0.69, indicating a good-to-strong cluster structure (Figure S6). Model features with the highest NMI scores (top 10 features) included cortical thickness of the pars triangularis, insula, middle temporal, supramarginal, superior and middle frontal gyrus regions and the SWAN inattention score (Table 2). The SWAN hyperactivity/impulsivity score and right pallidum volume were the only model features besides additional cortical thickness regions ranked among the top 35 features contributing to participant similarity. White matter FA measures did not prominently drive clustering (NMI scores were in the bottom half of all included features).

**Table 2.** The top 35 ranking model features contributing to participant similarity as determined by normalized mutual information and corresponding statistical summaries of between group differences for data-driven and diagnostic groups

| Rank | Top Contributing Features | Normalized<br>Mutual<br>Information | Data-driven Groups |  |  | Diagnostic Groups |  |  |
| --- | --- | --- | --- | --- | --- | --- | --- | --- |
|  |  |  | Statistic | pFDR | Eta<br>squared | Statistic | pFDR | Eta<br>squared |
| 1 | Right pars triangularis CT | 0.182 | 20.3 | 1.06E-09 | 0.315 | 0.33 | n.s | 0.0047 |
| 2 | Right insula CT | 0.182 | 24.9 | 4.05E-11 | 0.355 | 1.14 | n.s | 0.016 |
| 3 | SWAN Inattention Score | 0.181 | 16.0 | 3.46E-08 | 0.268 | 8.82 | 0.0086 | 0.11 |
| 4 | Right middle temporal CT | 0.180 | 28.2 | 3.01E-12 | 0.392 | 0.33 | n.s | 0.0046 |
| 5 | Left supramarginal CT | 0.172 | 23.7 | 9.67E-11 | 0.343 | 0.83 | n.s | 0.012 |
| 6 | Right inferior temporal CT | 0.169 | 17.5 | 9.36E-09 | 0.285 | 0.94 | n.s | 0.013 |
| 7 | Left middle temporal CT | 0.165 | 29.9 | 1.28E-12 | 0.404 | 1.92 | n.s | 0.027 |
| 8 | Right superior frontal CT | 0.165 | 19.7 | 1.60E-09 | 0.308 | 0.25 | n.s | 0.0036 |
| 9 | Left rostral middle frontal CT | 0.162 | 21.7 | 3.68E-10 | 0.321 | 0.85 | n.s | 0.012 |
| 10 | Left insula CT | 0.161 | 21.2 | 5.12E-10 | 0.307 | 1.43 | n.s | 0.019 |
| 11 | Left fusiform CT | 0.159 | 19.3 | 2E-09 | 0.308 | 2.84 | n.s | 0.039 |
| 12 | Left pars opercularis CT | 0.158 | 23.3 | 1.12E-10 | 0.350 | 0.63 | n.s | 0.0092 |
| 13 | Right inferior parietal CT | 0.158 | 20.2 | 1.09E-09 | 0.306 | 1.43 | n.s | 0.019 |
| 14 | Right rostral middle frontal CT | 0.157 | 18.0 | 6.33E-09 | 0.292 | 0.56 | n.s | 0.0084 |
| 15 | Left inferior parietal CT | 0.157 | 20.4 | 1.0E-09 | 0.304 | 0.94 | n.s | 0.013 |
| 16 | Left inferior temporal CT | 0.153 | 21.8 | 3.68E-10 | 0.335 | 1.63 | n.s | 0.024 |
| 17 | Right precentral CT | 0.149 | 13.2 | 6.66E-07 | 0.228 | 0.22 | n.s | 0.0031 |
| 18 | Left lateral orbitofrontal CT | 0.148 | 16.6 | 1.88E-08 | 0.261 | 0.69 | n.s | 0.0093 |
| 19 | Left lateral occipital CT | 0.148 | 15.1 | 8.89E-08 | 0.253 | 1.34 | n.s | 0.018 |
| 20 | Left lingual CT | 0.145 | 10.2 | 1.32E-05 | 0.181 | 1.41 | n.s | 0.018 |
| 21 | Right supramarginal CT | 0.145 | 17.2 | 1.24E-08 | 0.272 | 0.23 | n.s | 0.0031 |
| 22 | Left pars triangularis CT | 0.137 | 17.5 | 9.36E-09 | 0.281 | 0.22 | n.s | 0.0031 |
| 23 | Right fusiform CT | 0.136 | 16.9 | 1.43E-08 | 0.278 | 0.94 | n.s | 0.013 |
| 24 | Right lateral orbitofrontal CT | 0.134 | 13.1 | 6.94E-07 | 0.224 | 0.89 | n.s | 0.012 |
| 25 | Left postcentral CT | 0.134 | 15.2 | 7.8E-08 | 0.259 | 1.97 | n.s | 0.028 |
| 26 | SWAN Hyperactivity Score | 0.132 | 7.59 | 2.35E-04 | 0.146 | 6.14 | n.s | 0.083 |
| 27 | Right lateral occipital CT | 0.132 | 19.2 | 2.18E-09 | 0.299 | 1.99 | n.s | 0.027 |
| 28 | Left superior frontal CT | 0.131 | 13.9 | 2.91E-07 | 0.239 | 0.27 | n.s | 0.0039 |
| 29 | Right medial orbitofrontal CT | 0.127 | 12.9 | 8.3E-07 | 0.217 | 0.37 | n.s | 0.0051 |
| 30 | Left pericalcarine CT | 0.126 | 7.49 | 0.000258 | 0.143 | 0.21 | n.s | 0.0029 |
| 31 | Left caudal middle frontal CT | 0.123 | 19.6 | 1.69E-09 | 0.310 | 0.62 | n.s | 0.0093 |
| 32 | Right caudal middle frontal CT | 0.119 | 18.6 | 3.71E-09 | 0.296 | 0.1 | n.s | 0.0015 |
| 33 | Right postcentral CT | 0.118 | 10.4 | 1.01E-05 | 0.194 | 3.25 | n.s | 0.047 |
| 34 | Left pars orbitalis CT | 0.115 | 15.0 | 9.18E-08 | 0.253 | 0.45 | n.s | 0.0067 |
| 35 | Right pallidum | 0.115 | 8.02 | 0.000155 | 0.154 | 0.94 | n.s | 0.014 |

Group differences were determined using ANCOVAs while including age, sex and IQ as covariates.

**Figure 1.**
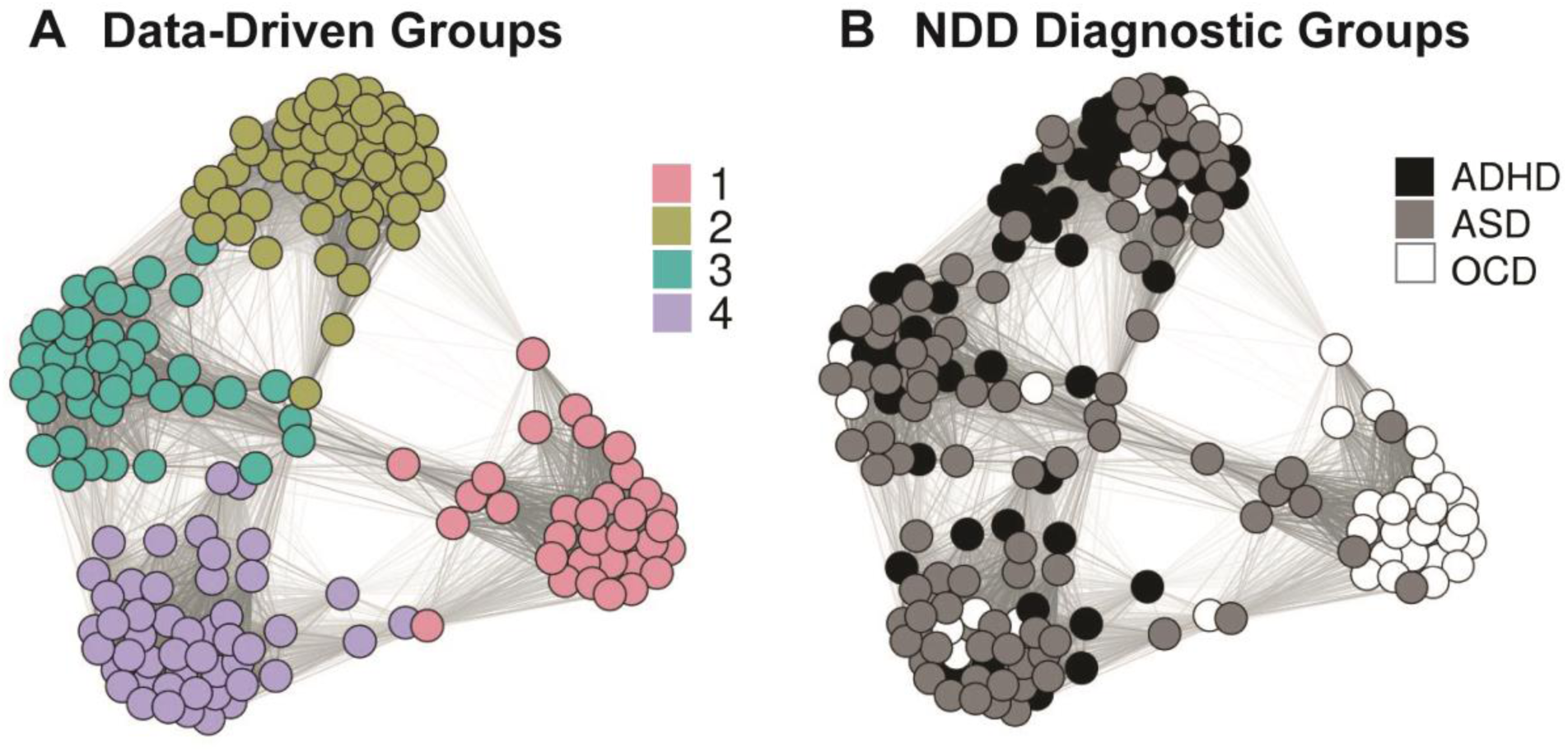
Representations of relative participant-participant similarities derived from the final SNF similarity matrix labelled by (A) data-driven group and by (B) diagnostic label.

### Influence of Demographic and IQ Variables on Data-Driven Groups

There was a significant effect of age on data-driven groups (F_3,172_=13.1, p<0.0001), due to younger age, on average, among Group 3 participants compared to all other groups (Figure S2). However, differences between groups on top ranking features were consistent across age (Figure 2A-C), and no group-by-age interaction effects were shown after FDR correction. There was also an effect of IQ (F_3,132_=4.5, p=0.005) and sex (X2=17.0, p=0.0007) on data-driven groups (Table S3), due to higher IQ in Group 1 compared to all other groups and proportionally more females in Groups 1 and 2 compared to Groups 3 and 4 (Figure S2).

**Figure 2.**
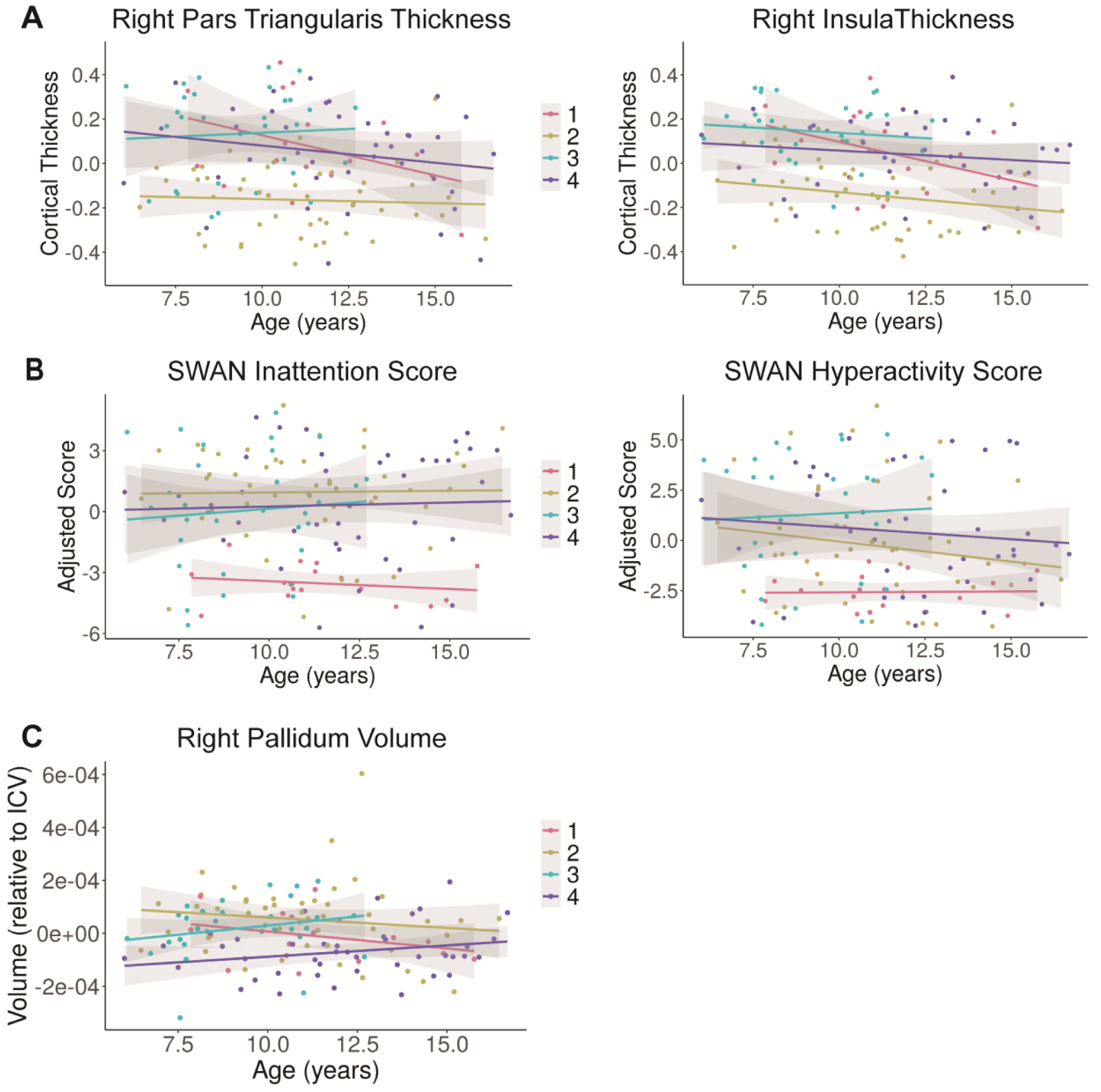
Figure panels depict features ranking among the top 35 features contributing to participant similarity, including top ranking (A) cortical thickness regions, (B) behavioural measures, and (C) subcortical volume plotted across age. Graphs show values after the effects of sex and IQ have been regressed out. Shaded areas represent 95% confidence intervals. Similar group difference patterns were found across other top contributing brain features within the same data type (Figure S1).

### Comparison of Top Contributing Features Between Data-Driven vs. NDD Groups

One-way ANCOVAs showed a significant main effect of group (FDR-corrected p<0.05) across top ranking model features (i.e. top 35), after controlling for the effects of age, sex and IQ (Table 2). With the exception of the SWAN inattention score (where effect size was much smaller), top ranking SNF features were not different between NDD groups. The top two model features with the largest effect sizes for data-driven group differences from each of the four data types examined included: bilateral middle temporal thickness (left: F_3,129_=29.9, p=1.28e-12, η2=0.40; right: F_3,129_=28.2, p=3.01e-12, η2=0.39), right thalamus (F_3,129_=12.6, p=1.04e-6, η2=0.23), right nucleus accumbens volume (F_3,129_=12.8, p=8.76e-07, η2=0.22), FA in the right posterior limb of the internal capsule (F_3,129_=3.73, p=0.02, η2=0.08), right cingulum FA (F_3,129_=3.2, p=0.04, η2=0.06), SWAN inattention score (F_3,129_=16.0, p=3.46e-8, η2=0.27), SCQ score (F_3,129_=11.4, p=3.75e-6, η2=0.19)(Table S3). In contrast, there were no significant differences between NDD groups on any of the brain features included in the SNF analysis; effect sizes were typically smaller by ten fold (or more) (Table 2, Table S3).

### Brain-Behavior Profiles of Data-Driven Groups Based on Top Model Features

Follow-up Tukey comparison tests (Table S3) examining group differences on top contributing model features suggested that Groups 2, 3, and 4 had greater impairments on brain and behavioral measures (compared to Group 1), although specific profiles differed. Group 2 (n=54), consisted of a near even split of children with ADHD (n=24) or ASD (n=23), and fewer OCD (n=7). Group 2 consistently featured significant decreases in cortical thickness within regions with top NMI scores (e.g. right pars triangularis) compared to Groups 1, 3 and 4 (Figure S4). In contrast, Group 3 (n=41) had the highest cortical thickness relative to other groups, was comprised mainly of children with ASD (n=25), a sizeable minority with ADHD (n=13), and few with OCD (n=3), and was male dominant. Group 4 (n=48) included predominantly children with ASD (n=31), with a sizeable minority of ADHD (n=12), and fewer children with OCD (n=5), and was male dominant. Group 4 featured significant decreases in right pallidum volume (the top NMI subcortical measure) compared to Groups 2 and 3. Groups 3 and 4 were the most behaviorally impaired. SWAN inattention and hyperactivity/impulsivity scores were significantly lower in Group 1 (n=33), which was comprised mainly of children with OCD (n=24) and fewer with ASD (n=9).

Notably the cortical thickness and subcortical measures in Group 1 were neither highest nor lowest. Internal capsule and cingulum FA (white matter regions with largest between group effects) were increased in Group 4 versus Group 2.

### Cluster Evaluation

#### Cluster Stability

Top ranking features based on NMI score remained consistent across resampling. Of the top 10 features, 8 remained within the top 35 ranking features across 60-90% of clustering permutations, including SWAN inattention score; the remaining 2 features originally among the top 10 (right inferior temporal and left supramarginal thickness) remained within the top 35 across 40-42% of permutations. SWAN hyperactivity/impulsivity score and right pallidum volume remained among the top 35 features across 68% and 52% of permutations, respectively. Other subcortical volumes were also among the top 35 features across a substantial number of permutations (e.g. right thalamus: 65%). Across resampling, any given participant clustered with each other participant within their group on average 67% of the time, as compared to across them (9.4% of the time, Figure S5). An Adjusted Rand Index of 0.46 across 1000 iterations was found, indicating over 70% agreement across clusters[39].

### Extension of Data-Driven Group Distinctions to Out-of-Model Features

In the three out of model ‘phenotypes’ [adaptive (everyday) functioning, surface area, brain structural covariance network indices], we found that effect sizes for differences between data-driven groups were larger than for NDD groups. For data-driven groups, the effect size of the between-group difference on the General Adaptive Composite score (F3,126=9.1, p=1.8E-5, η2=0.16) was larger compared to NDD groups (F2,127=8.1, p=4.9E-4, η2=0.10) (see Figure 3), when covarying for sex, age and IQ. Surface area was generally not different among either data-driven or NDD groups. For structural covariance network indices, effects were larger among data-driven compared to NDD groups for network strength and density across thresholds (See Supplementary Materials and Methods for detailed results).

**Figure 3.**
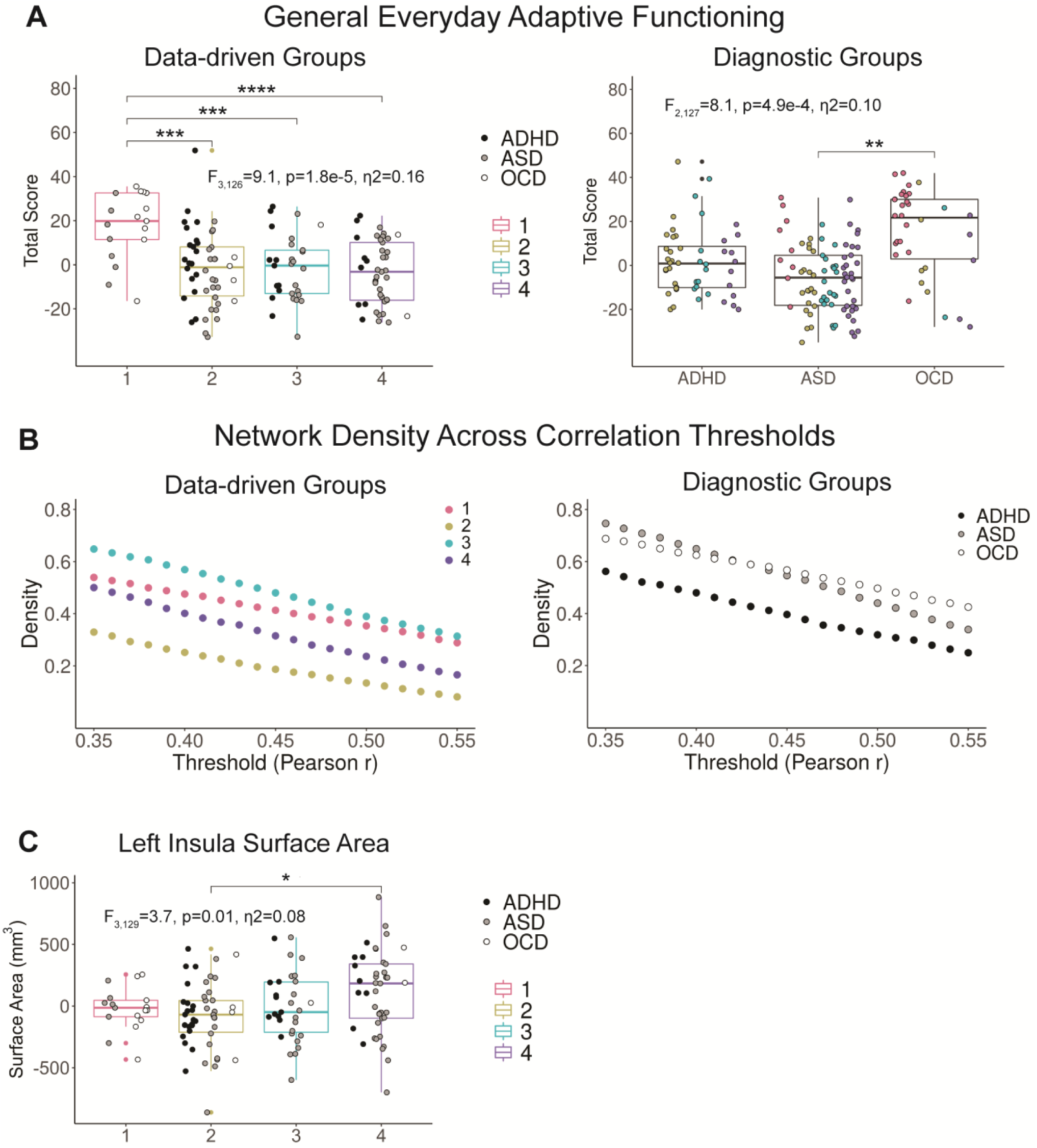
(A) ABAS-II General Adaptive Composite scores for both data-driven and diagnostic groups after regressing out the effects of age, sex and IQ. (B) Structural covariance network densities across a range of Pearson r thresholds for both data-driven and diagnostic groups. Networks were created from cortical thickness regions after effects of age, sex and IQ were regressed out. (C) Left insula surface area in data-driven groups after regressing out the effects of age, sex and IQ (**** =p<0.0001, ***= p<0.001, **= p<0.01, *=p <0.05). Boxplots show the first and third quartile with extensions representing 1.5 * the inter-quartile range.

#### Comparison of Classification Accuracy Based on Different Selections of Top Ranking Features

Model performance was highest when the top 2 ranking features (highest NMI scores) from each of the four data types (i.e., right pars triangularis, right insula thickness, SWAN inattention and hyperactivity/impulsivity, right pallidum, right putamen volume, left anterior limb internal capsule FA, right retrolenticular internal capsule FA), or all 135 input features were included in the classifier (Figure S7), compared to when only top ranking features were included in the classifier. Mean sensitivity performance for detecting data-driven groups when including the top 2 features across data types in the classifier ranged from 62-75%, exceeding chance. Mean specificity performance was >80%. When the top 10 or 35 ranking features were included in the classifier, performance remained stable for prediction of Groups 1 and 2, but comparatively declined for Groups 3 and 4. When features from only one of the four data types were included, mean sensitivity dropped to below 50% for at least two of Groups 1, 3 and 4.

## DISCUSSION

By fusing across multiple brain imaging phenotypes and behavioral measures, we identified novel transdiagnostic data-driven groups, which feature more homogeneous characteristics within groups in both brain and behavioral measures compared to current DSM-5 categories (ASD, ADHD, OCD). In particular, we found that cortical thickness in regions important for social behavior (inferior frontal gyrus, insula, inferior parietal cortex, temporal cortex) and executive function (superior and middle frontal gyrus) along with inattention scores were the top contributors to the model. These differences were consistent across the age range, a period of dynamic brain growth and change[40,41]. Data-driven participant similarity groups displayed internal stability of clustering, and stability of top contributing model features and pairwise participant clustering across resampling. Stronger differences between data-driven groups compared to DSM-5 diagnostic groups extended to clinically and biologically relevant features excluded from the SNF analysis, most notably adaptive (everyday) functioning. Although white matter FA was not among the top contributing features, classification accuracy was best when this data type was included in the model.

Of the four data-driven groups, Groups, 2, 3, and 4 were the most behaviorally and functionally impaired, consistent with alterations in brain imaging measures. However, among these groups, there were notable differences, suggesting different neurobiological features may relate to different behavioral profiles. Group 2, comprised evenly of children with ADHD or ASD, had higher inattention scores and a distinct pattern of decreased cortical thickness compared to all other groups. The Group 2 profile found in the current study may be most consistent with prior work showing delayed and deviated cortical and neural network maturation, particularly in frontal regions in large-scale studies of children with ADHD, including reduced thickness[42,43]. In direct contrast to Group 2, Group 3 showed elevated hyperactivity and higher cortical thickness in top ranking regions, but was also the youngest group and in earlier stages of normative behavioural and cortical development, perhaps accounting for some of these differences[40]. Nevertheless, plotting cortical thickness across age showed that these findings were sustained and age-independent. These contrasting cortical thickness phenotypes were not elicited through diagnostic comparisons (for which there were no significant differences in thickness). By contrasting data-driven versus diagnostic groups and via inclusion of multiple imaging phenotypes (i.e., cortical thickness, subcortical volumes, white matter FA) our findings build on a previous analysis of the POND sample[22]. In addition, the novelty of our findings is also notable because of out-of-model differences among the data-driven groups in adaptive functioning and brain network structure. Similar frontal and temporal cortical regions have also been implicated in a mega-analysis from the ENIGMA group comparing ASD to typically developing controls, with both increased and decreased thickness found in ASD[44]. Our work suggests that these same regions contribute to biological differences among data-driven groups. It is possible that reduced cortical thickness in some cases versus increased thickness in others (which may reflect delayed maturation) exists in subgroups of children with the same NDD diagnosis, but is associated with different behavioral phenotypes (e.g., more inattention vs. more hyperactivity).

Group 4 was characterized biologically by decreases in striatal and thalamic subcortical volumes. Reductions in similar regions were shown in ASD in the recent case-control ENIGMA ASD mega-analysis[44]. When this group of features are taken together, involvement of the cortico-striatal-thalamic-cortical (CSTC) circuit also emerges as a notable pattern. The CSTC is a network widely implicated as vulnerable in ASD, ADHD and OCD[45–49]. Children in this group may represent those with this shared vulnerability pathway cutting across diagnoses.

In contrast to Groups 2-4, Group 1 was largely comprised of children with OCD. This group featured reduced impairment across included behavioural features (except for OCD symptoms), lacked distinguishable biological impairments, and had higher IQ scores than Groups 3 and 4. Some prior studies using POND data have found, on average, children with OCD have milder impairments at the brain and behavioural level compared to children with ASD or ADHD[12,22,35]. Our results identified some children with ASD that also feature milder impairments and fit into this group. Group 1 also featured highest adaptive functioning compared to Groups 2, 3 and 4 on out-of-model evaluation. Longitudinal analyses are needed to track whether outcomes differ in this group compared to other groups over time. If lower adaptive functioning impairment in this group remains stable over time, features differentiating this group could potentially have clinical utility for identifying children with NDDs that may have favourable outcomes or respond differently to available treatment or clinical management approaches.

Although modest, brain network comparisons among the groups provided support for generalizability of distinctions between data-driven groups to features that were not utilized to delineate groups. In particular, the lower network density in Group 2 among cortical thickness regions supported the impaired cortical thickness phenotype found in this group using the data-driven model. Lower network density in Group 2 (and 4) may indicate broader, more wide-ranging network based alterations associated with their respective behavioural alteration profiles, and perhaps earlier developmental insults affecting more of the brain[50].

It is notable that females with a diagnosis of OCD or ADHD mainly clustered into Groups 1 and 2, while females with ASD clustered across data-driven groupings. Biological sex is an important source of heterogeneity in NDDs[11] and aspects of sex-specific brain structure and functional connectivity patterns have been found in ASD[51] and to a lesser degree in ADHD[52] or OCD[53]. Previous evidence has suggested a protective effect for females, or required increased biological ‘hit’ related to resilience to developing NDDs[54,55]. However, interpretation of any differences found amongst males versus females in the current study is limited due to the small numbers of represented females with NDDs.

We took a series of approaches to determine the stability of data-driven grouping and potential meaningfulness. We found that top contributing features could reliably identify our groups (mean sensitivity 42-86%), which may improve with larger sample sizes. Although all groups were identifiable using the full spectrum of features included in our SNF analysis, sensitivity for prediction of Groups 1 and 2 remained stable when based on a more constrained set of top contributing features (i.e. cortical thickness and inattention/hyperactivity-impulsivity symptom scores), suggesting that these groups may be identifiable in another sample using more constrained behavioral and biological information. In contrast, a fuller spectrum of data may be needed to identify children with more complex presentations (and perhaps more overlapping behavioral and functional impairment profiles) as in Groups 3 and 4. Although Group 1 was the most stable across resampling, participants in this group were not classified with the highest sensitivity, perhaps due to their intermediate values on biological measures.

### Limitations

Results should be interpreted in the context of study limitations. Sample sizes were unequal across diagnostic groups, and larger numbers could have provided a number of statistical advantages. Future work could extend the dimensionality of input features to include cognitive, genetic, environmental and other neuroimaging features. Visualizing similarities between participants showed that although participants within the four data-driven groups identified featured more similar and separable brain-behaviour profiles than found using conventional DSM-5 diagnostic categories, some participants did not cluster ‘cleanly’ into a specific data-driven group. Although studying individuals on a spectrum may be more informative for characterizing the continuum of brain-behavioural relationships present across the population (and shown to be relevant on clustering of children with different NDDs[22], others argue that biotypes (i.e. new subgroups) may be needed to parse multi-dimensional brain-behaviour profiles into groupings that can be useful for clinical translation[10].

## Conclusion

The current study adds to recent work suggesting biological and behavioral convergences across NDDs, as well as divergences within them [16,22,56]. We identified new groups cutting across NDDs characterized by multi-level neuroimaging and behavioral data. The more similar biological profiles found among our data-driven groups invites future work to replicate findings, test longitudinally for prognostic value as well as stratification for targeted treatment studies.

## ACKNOWLEDGEMENTS

We thank the following individuals for research support and data collection: Tara Goodale, M.Sc., Reva Schachter, M.Sc., Mithula Sriskandarajah, B.Sc., Marlena Colasanto, M.Sc., Jennifer Gomez, M.A., and Laura Park, M.Sc, from The Hospital for Sick Children; Susan Day Fragiadakis, M.A., Naomi Peleg, M.Sc.,and Leanne Ristic, B.A., from Holland Bloorview Kids Rehabilitation Hospital; Richa Mehta, B.A., Christina Sommerdyk, M.Sc., from the Lawson Health Research Institute; Carolyn Russell,B.Sc., Alessia Greco, M.A., Mike Chalupka, B.A., B.Sc., Christina Chrysler, B.A., Irene O’Connor, M.Ed. Psych., from McMaster Children’s Hospital. We thank Margot J. Taylor for her review of a draft version of the manuscript.

## REFERENCES

1. Szatmari P, Georgiades S, Duku E, Bennett TA, Bryson S, Fombonne E, et al. Developmental trajectories of symptom severity and adaptive functioning in an inception cohort of preschool children with autism spectrum disorder. JAMA Psychiatry. 2015;72: 276–283.

2. Karalunas SL, Fair D, Musser ED, Aykes K, Iyer SP, Nigg JT. Subtyping attention-deficit/hyperactivity disorder using temperament dimensions: Toward biologically based nosologic criteria. JAMA Psychiatry. 2014;71: 1015–1024.

3. Di Rezze B, Duku E, Szatmari P, Volden J, Georgiades S, Zwaigenbaum L, et al. Examining Trajectories of Daily Living Skills over the Preschool Years for Children with Autism Spectrum Disorder. J Autism Dev Disord. 2019;49: 4390–4399.

4. Schwartzman CM, Boisseau CL, Sibrava NJ, Mancebo MC, Eisen JL, Rasmussen SA. Symptom subtype and quality of life in obsessive-compulsive disorder. Psychiatry Res. 2017;249: 307–310.

5. Anholt GE, Cath DC, Van Oppen P, Eikelenboom M, Smit JH, Van Megen H, et al. Autism and adhd symptoms in patients with ocd: Are they associated with specific oc symptom dimensions or oc symptom severity. J Autism Dev Disord. 2010. doi: 10.1007/s10803-009-0922-1

6. Ashwood KL, Tye C, Azadi B, Cartwright S, Asherson P, Bolton P. Brief report: Adaptive functioning in children with ASD, ADHD and ASD+ ADHD. J Autism Dev Disord. 2015;45: 2235–2242.

7. Van Der Meer JMJ, Oerlemans AM, Van Steijn DJ, Lappenschaar MGA, De Sonneville LMJ, Buitelaar JK, et al. Are autism spectrum disorder and attention-deficit/hyperactivity disorder different manifestations of one overarching disorder? Cognitive and symptom evidence from a clinical and population-based sample. J Am Acad Child Adolesc Psychiatry. 2012. doi: 10.1016/j.jaac.2012.08.024

8. Lionel AC, Crosbie J, Barbosa N, Goodale T, Thiruvahindrapuram B, Rickaby J, et al. Rare Copy Number Variation Discovery and Cross-Disorder Comparisons Identify Risk Genes for ADHD. Sci Transl Med. 2011. doi: 10.1126/scitranslmed.3002464

9. Constantino JN, Charman T. Diagnosis of autism spectrum disorder: reconciling the syndrome, its diverse origins, and variation in expression. Lancet Neurol. 2016;15: 279–291.

10. Feczko E, Miranda-Dominguez O, Marr M, Graham AM, Nigg JT, Fair DA. The Heterogeneity Problem: Approaches to Identify Psychiatric Subtypes. Trends Cogn Sci. 2019;23: 584–601.

11. Lombardo MV, Lai M-C, Baron-Cohen S. Big data approaches to decomposing heterogeneity across the autism spectrum. Mol Psychiatry. 2019. doi: 10.1038/s41380-018-0321-0

12. Ameis SH, Lerch JP, Taylor MJ, Lee W, Viviano JD, Pipitone J, et al. A Diffusion Tensor Imaging Study in Children With ADHD, Autism Spectrum Disorder, OCD, and Matched Controls: Distinct and Non-Distinct White Matter Disruption and Dimensional Brain-Behavior Relationships. Am J Psychiatry. 2016. doi: 10.1176/appi.ajp.2016.15111435

13. Dajani DR, Burrows CA, Odriozola P, Baez A, Beth M, Mostofsky SH, et al. NeuroImage : Clinical Investigating functional brain network integrity using a traditional and novel categorical scheme for neurodevelopmental disorders. NeuroImage: Clinical. 2019;21: 101678.

14. Di Martino A, Zuo X-N, Kelly C, Grzadzinski R, Mennes M, Schvarcz A, et al. Shared and distinct intrinsic functional network centrality in autism and attention-deficit/hyperactivity disorder. Biol Psychiatry. 2013;74: 623–632.

15. Chantiluke K, Christakou A, Murphy CM, Giampietro V, Daly EM, Ecker C, et al. Disorder-specific functional abnormalities during temporal discounting in youth with Attention Deficit Hyperactivity Disorder (ADHD), Autism and comorbid ADHD and Autism. Psychiatry Res. 2014;223: 113–120.

16. Boedhoe PSW, van Rooij D, Hoogman M, Twisk JWR, Schmaal L, Abe Y, et al. Subcortical brain volume, regional cortical thickness and cortical surface area across attention-deficit/hyperactivity disorder (ADHD), autism spectrum disorder (ASD), and obsessive-compulsive disorder (OCD). bioRxiv. 2019. p. 673012. doi: 10.1101/673012

17. Baribeau DA, Dupuis A, Paton TA, Hammill C, Scherer SW, Schachar RJ, et al. Structural neuroimaging correlates of social deficits are similar in autism spectrum disorder and attention-deficit/hyperactivity disorder: analysis from the POND Network. Transl Psychiatry. 2019;9: 72.

18. Aoki Y, Yoncheva YN, Chen B, Nath T, Sharp D, Lazar M, et al. Association of White Matter Structure With Autism Spectrum Disorder and Attention-Deficit/Hyperactivity Disorder. JAMA Psychiatry. 2017;74: 1120–1128.

19. Feczko E, Balba NM, Miranda-Dominguez O, Cordova M, Karalunas SL, Irwin L, et al. Subtyping cognitive profiles in Autism Spectrum Disorder using a Functional Random Forest algorithm. Neuroimage. 2018;172: 674–688.

20. Stefanik L, Erdman L, Ameis SH, Foussias G, Mulsant BH, Behdinan T, et al. Brain-Behavior Participant Similarity Networks Among Youth and Emerging Adults with Schizophrenia Spectrum, Autism Spectrum, or Bipolar Disorder and Matched Controls. Neuropsychopharmacology. 2017;43: 1180.

21. Hawco C, Buchanan RW, Calarco N, Mulsant BH, Viviano JD, Dickie EW, et al. Separable and Replicable Neural Strategies During Social Brain Function in People With and Without Severe Mental Illness. Am J Psychiatry. 2019;176: 521–530.

22. Kushki A, Anagnostou E, Hammill C, Duez P, Brian J, Iaboni A, et al. Examining overlap and homogeneity in ASD, ADHD, and OCD: a data-driven, diagnosis-agnostic approach. Transl Psychiatry. 2019;9: 318.

23. Wang B, Mezlini AM, Demir F, Fiume M, Tu Z, Brudno M, et al. Similarity network fusion for aggregating data types on a genomic scale. Nat Methods. 2014;11: 333.

24. Lord C, Rutter M, Le Couteur A. Autism Diagnostic Interview-Revised: a revised version of a diagnostic interview for caregivers of individuals with possible pervasive developmental disorders. J Autism Dev Disord. 1994;24: 659–685.

25. Lord C, Risi S, Lambrecht L, Cook EH, Leventhal BL, DiLavore PC, et al. The Autism Diagnostic Observation Schedule—Generic: A Standard Measure of Social and Communication Deficits Associated with the Spectrum of Autism. J Autism Dev Disord. 2000;30: 205–223.

26. Ickowicz A, Schachar RJ, Sugarman R, Chen SX, Millette C, Cook L. The parent interview for child symptoms: a situation-specific clinical research interview for attention-deficit hyperactivity and related disorders. Can J Psychiatry. 2006;51: 325–328.

27. Kaufman J, Birmaher B, Brent D, Rao U, Flynn C, Moreci P, et al. Schedule for Affective Disorders and Schizophrenia for School-Age Children-Present and Lifetime Version (K-SADS-PL): initial reliability and validity data. J Am Acad Child Adolesc Psychiatry. 1997;36: 980–988.

28. Scahill L, Riddle MA, McSwiggin-Hardin M, Ort SI, King RA, Goodman WK, et al. Children’s Yale-Brown Obsessive Compulsive Scale: reliability and validity. J Am Acad Child Adolesc Psychiatry. 1997;36: 844–852.

29. Achenbach TM, Edelbrock CS, Others. Manual for the child behavior checklist and revised child behavior profile. 1983. Available: https://pdfs.semanticscholar.org/e06f/18f950ee20811acd25b9671c14a80b681e3c.pdf

30. Park LS, Burton CL, Dupuis A, Shan J, Storch EA, Crosbie J, et al. The Toronto Obsessive-Compulsive Scale: Psychometrics of a Dimensional Measure of Obsessive-Compulsive Traits. J Am Acad Child Adolesc Psychiatry. 2016;55: 310–318.e4.

31. Berument SK, Rutter M, Lord C, Pickles A, Bailey A. Autism screening questionnaire: diagnostic validity. Br J Psychiatry. 1999;175: 444–451.

32. Bodfish JW, Symons FJ, Parker DE, Lewis MH. Varieties of repetitive behavior in autism: comparisons to mental retardation. J Autism Dev Disord. 2000;30: 237–243.

33. Swanson J, Schuck S, Mann M, Carlson C, Hartman K, Sergeant J, et al. Categorical and dimensional definitions and evaluations of symptoms of ADHD: The SNAP and SWAN rating scales. University of California, Irvine. 2006.

34. Harrison PL, Oakland T. Adaptive behavior assessment system. 2003.

35. Baribeau DA, Doyle-Thomas KAR, Dupuis A, Iaboni A, Crosbie J, McGinn H, et al. Examining and comparing social perception abilities across childhood-onset neurodevelopmental disorders. J Am Acad Child Adolesc Psychiatry. 2015;54: 479–86.e1.

36. Fischl B. FreeSurfer. Neuroimage. 2012;62: 774–781.

37. Smith SM, Jenkinson M, Johansen-Berg H, Rueckert D, Nichols TE, Mackay CE, et al. Tract-based spatial statistics: voxelwise analysis of multi-subject diffusion data. Neuroimage. 2006;31: 1487–1505.

38. Johns Hopkins University. School of Medicine, Jean S. The Johns Hopkins Atlas of Human Functional Anatomy. JHU Press; 1997.

39. Steinley D, Brusco MJ, Hubert L. The variance of the adjusted Rand index. Psychol Methods. 2016;21: 261–272.

40. Sussman D, Leung RC, Chakravarty MM, Lerch JP, Taylor MJ. Developing human brain: Age-related changes in cortical, subcortical, and cerebellar anatomy. Brain Behav. 2016. doi: 10.1002/brb3.457

41. Oyefiade AA, Ameis S, Lerch JP, Rockel C, Szulc KU, Scantlebury N, et al. Development of short-range white matter in healthy children and adolescents. Hum Brain Mapp. 2018;39: 204–217.

42. Shaw P, Greenstein D, Sharp W, Clasen L, Giedd J, Rapoport J, et al. Longitudinal mapping of cortical thickness and clinical outcome in children and adolescents with attention-deficit/hyperactivity disorder. Arch Gen Psychiatry. 2006;63: 540–549.

43. Shaw P, Eckstrand K, Sharp W, Blumenthal J, Lerch JP, Greenstein D, et al. Attention-deficit/hyperactivity disorder is characterized by a delay in cortical maturation. Proceedings of the National Academy of Sciences. 2007;104: 19649–19654.

44. Van Rooij D, Anagnostou E, Arango C, Auzias G, Behrmann M, Busatto GF, et al. Cortical and subcortical brain morphometry differences between patients with autism spectrum disorder and healthy individuals across the lifespan: Results from the ENIGMA ASD working group. Am J Psychiatry. 2018;175: 359–369.

45. Zhu Y, Jiang X, Ji W. The Mechanism of Cortico-Striato-Thalamo-Cortical Neurocircuitry in Response Inhibition and Emotional Responding in Attention Deficit Hyperactivity Disorder with Comorbid Disruptive Behavior Disorder. Neurosci Bull. 2018;34: 566–572.

46. Kuo H-Y, Liu F-C. Synaptic Wiring of Corticostriatal Circuits in Basal Ganglia: Insights into the Pathogenesis of Neuropsychiatric Disorders. eNeuro. 2019;6. doi: 10.1523/ENEURO.0076-19.2019

47. Fettes P, Schulze L, Downar J. Cortico-Striatal-Thalamic Loop Circuits of the Orbitofrontal Cortex : Promising Therapeutic Targets in Psychiatric Illness. 2017;11: 1–23.

48. Hoogman M, Bralten J, Hibar DP, Mennes M, Zwiers MP, Schweren LSJ, et al. Subcortical brain volume differences in participants with attention deficit hyperactivity disorder in children and adults: a cross-sectional mega-analysis. Lancet Psychiatry. 2017;4: 310–319.

49. Boedhoe PSW, Schmaal L, Abe Y, Ameis SH, Arnold PD, Batistuzzo MC, et al. Distinct Subcortical Volume Alterations in Pediatric and Adult OCD: A Worldwide Meta- and Mega-Analysis. Am J Psychiatry. 2017;174: 60–69.

50. Bethlehem RAI, Romero-Garcia R, Mak E, Bullmore ET, Baron-Cohen S, Bethlehem RAI. Structural Covariance Networks in Children with Autism or ADHD. Cereb Cortex. 2017;27: 4267–4276.

51. Lai M-C, Lerch JP, Floris DL, Ruigrok ANV, Pohl A, Lombardo MV, et al. Imaging sex/gender and autism in the brain: Etiological implications. J Neurosci Res. 2017;95: 380–397.

52. Scofield JE, Johnson JD, Wood PK, Geary DC. Latent resting-state network dynamics in boys and girls with attention-deficit/hyperactivity disorder. PLoS One. 2019;14: e0218891.

53. Hawco C, Voineskos AN, Radhu N, Rotenberg D, Ameis S, Backhouse FA, et al. Age and gender interactions in white matter of schizophrenia and obsessive compulsive disorder compared to non-psychiatric controls: commonalities across disorders. Brain Imaging Behav. 2017;11: 1836–1848.

54. Werling DM. The role of sex-differential biology in risk for autism spectrum disorder. Biol Sex Differ. 2016;7: 58.

55. Taylor MJ, Lichtenstein P, Larsson H, Anckarsäter H, Greven CU, Ronald A. Is There a Female Protective Effect Against Attention-Deficit/Hyperactivity Disorder? Evidence From Two Representative Twin Samples. J Am Acad Child Adolesc Psychiatry. 2016;55: 504–512.e2.

56. Brem S, Grünblatt E, Drechsler R, Riederer P, Walitza S. The neurobiological link between OCD and ADHD. Atten Defic Hyperact Disord. 2014;6: 175–202.

