## Supplemental Information for "Integration of Brain and Behavior Measures for Identification of Data-Driven Groups Cutting Across Children with ASD, ADHD, or OCD"

### MRI Acquisition

T1-weighted scans were acquired using a MPRAGE sequence (0.8x0.8x0.8 mm3, TR=1870ms, TE=3.14ms, TI=945ms, Flip Angle=9°, sagittal FOV=256x240mm2, 240 Slices, GRAPPA=2). Single shell diffusion image acquisitions were acquired in three blocks (sequence: (2x2x2 mm3, TR=3800ms, TE=73ms, Flip Angle=90° FOV=244x244mm2, 70 Slices) which were concatenated together for processing. Acquisition was divided over 3 consecutive scans with 19, 20 or 21 gradient directions along with 3 B0's acquired in each. Total scan time was 11:27 (3:40 + 3:49 + 3:58) minutes.

### Diffusion Weighting Imaging Preprocessing

Diffusion MRI data from the three runs were concatenated. If multiple run data was acquired the data was visually examined and the better run selected. Data were next denoised and upsampled (to 1x1x1) using the MRtrix3's dwidenoise and upsample commands respectively (http://www.mrtrix.org/). Images were then corrected for motion artifacts while simultaneously accounting for field inhomogeneities (by way of fieldmap correction) and eddy current induced artifacts using FSL's eddy function (Andersson and Sotiropoulos 2016; Andersson et al. 2016).

### MRI Quality Control

### T1-weighted MRI

Participant exclusion based on motion related artifacts or poor scan quality was carried out using a step-wise QC protocol visually by a trained rater and quantitatively using image quality metrics. Visual QC consisted of inspecting brain images pre-processed through fMRIprep (Esteban et al. 2019). A stepwise rating system was followed and each scan was given a score of 1-5 depending on the severity of the artifacts and their influence on segmentation, with 1 being the least affected by artifacts and 5 being the most (similar to (Pardoe, Kucharsky Hiess, and

Kuzniecky 2016)). From there, the ratings were grouped into pass (rating of 1-3) and fail (rating of 4-5). To reduce some of the confounds of a purely visual rating system, we also evaluated three important quality metrics associated with noise derived from MRIQC (Esteban et al. 2017). These were: contrast-to-noise ratio (CNR), signal-to-noise ratio (SNR) and coefficient of joint variation (CJV). SNR is conceptualized as the signal of the of the MR image compared to the background noise. A higher ratio is more desirable (Dietrich et al. 2007). CNR is an extension of SNR and determines how separated the tissue distributions between white and gray matter are. A higher ratio is more desirable (Magnotta et al. 2006). CJV is a proposed function to determine optimization of intensity non-uniformity in a brain image, with higher values associated with increased head motion and more noise (Ganzetti et al. 2016). Participants were removed based on the visual QC, but also excluded if their values of CNR, SNR, and/or CJV exceeded 2 standard deviations of the sample.

**Diffusion-weighted MRI**

Similarly, for diffusion scans, participants were excluded based on visual inspection for motion related artifacts or poor scan quality carried out using a step-wise QC protocol by a trained rater. FSL's DTIFIT was run to fit a diffusion tensor model to each voxel and produce output to assess slices from each orthogonal view of the B0, brain mask, first eigenvector overlaid onto the fractional anisotropy map, and sum of squared errors. Participants were removed based on evidence for slice dropout, inter-slice variability, and truncated brain regions.

**Demographics of excluded participants**

A total of 90 participants were excluded from analyses (Figure S1). There were significant differences between included and excluded children and youth in clinical diagnoses ($\chi_2$=20.9, df=2, p<0.0001). Most excluded participants had a clinical diagnosis of ASD (n=71), followed by ADHD (n=12) and OCD (n=7). Excluded participants were slightly younger (mean age = 10.2, SD=4.5) compared to those included (mean age = 11.2, SD=2.6) ($t_{1,121}$=2.1, p=0.04). Excluded participants also had lower IQ (mean=93.5, SD=20.6) compared to included participants (mean=101.1, SD=17.1) ($t_{1,65}$=2.2, p=0.03). There was no difference in sex between included and excluded participants ($X_2$=0.6, df=1, p=0.4).

**Medications**

Of the 176 children and youth included in this study, 60 (34%) of them were currently on or had been on a prescription medication in the past 6 months. A total of 59 (34%) reported that they were not taking any medication and there was no information on medications prescribed for 57 (32%) participants in the sample. Among participants diagnosed with ASD with information, 22/42 were taking medications. Among participants diagnosed with ADHD, 24/49 were taking medications, and with OCD 14/38 were on medications. No significant between group difference in medication status was found between data-driven groups ($X_2$=4.6, df=6, p=0.6). For data-driven groups, the portion of participants with information on medication treatment indicated that in Group 1:11/19, Group 2: 21/41, Group 3: 14/29, and in Group 4: 14/30 participants were taking medications.

**SNF Data Integration and Clustering Analysis**

SNF (v2.3.0 R package) was used to integrate multimodal data types. This method works by first creating separate participant similarity networks for each data type followed by a nonlinear combination method based on message-passing theory that iteratively fuses data types into a single participant similarity network (Wang et al. 2014). Similarity matrices for each of the four data types (cortical thickness, subcortical volume, fractional anisotropy & behavioural measures) were calculated using Euclidean distance with a nearest neighbours value of 18, normalization parameter of 0.8, and iteration number of 10. See Table S3 for a list of all included features. Normalized mutual information (NMI) was used to measure the overlap in a similarity matrix created using any single feature included in the model within each data type and the similarity matrix defined by all of the data combined.

Nearest neighbours (K) and normalization (alpha) parameters were chosen in consultation with developers to increase NMI scores in accordance with the recommended ranges (K=10-30 and alpha=0.3-0.8) and a suggested nearest neighbours value of total sample/10=18. Cluster number was chosen using the SNF *estimateNumberOfClustersGivenGraph* function, which evaluates eigen-gaps and rotation cost. To determine stability of suggested cluster number, this function was run on a series of similarity networks determined using the range of nearest neighbours and normalization parameters noted above. Four clusters was consistently optimal based on both eigen-gaps and rotation cost. NMI scores and the number of optimal clusters were also assessed across the recommended range of number of iterations between 10-20. Neither cluster number nor NMI were found to vary much across this parameter, and thus 10 was chosen for the number of iterations.

Spectral clustering (SNF *spectralClustering* function) was then applied to identify subgroups from a consensus fused similarity matrix defined based on all 135 features across 1000 iterations of resampling 80% of the participants. Silhouette plots derived from the consensus matrix were used to quantify similarity for participants within a given cluster compared to participants in all other clusters (Figure S6).

**Cluster Stability**

To evaluate reproducibility, we conducted resampling-based stability testing with 1000 iterations. In each iteration, 80% of participants were randomly sampled proportional to group size, their features integrated using SNF and then clustered into subgroups. For each iteration, the following were calculated: 1) the consistency of the top 10 and 35 contributing features based on NMI scores, 2) the percentage of time that each participant clustered with every other participant (when participants were both included in that resampling iteration)(Figure S5), and 3) the overlap between cluster partitions measured using the Adjusted Rand Index (this index ranges between 0, indicating completely random cluster overlap, and 1, indicating identical clustering across iterations) (Hubert and Arabie 1985).

**Out-of-model Features - Methods**

Data-driven group and diagnostic group differences were examined on brain and behaviour features excluded from SNF analysis, including: 1) ABAS-II General Adaptive Composite Score, 2) surface area within the regions corresponding to the top 10 cortical thickness features based on NMI, and 3) global efficiency, network strength, and network density of cortical thickness structural covariance networks across a range of thresholds.

Structural covariance networks were created by first regressing out age, sex and IQ from cortical thickness values. Then, within each group, between-subject Pearson correlation coefficients (r) were calculated for each pairwise combination of brain regions to create group-wide structural covariance matrices for each data-driven and NDD (diagnostic) group. A range of density thresholds at 5% intervals between 10 and 50% were applied to edges and binarized to produce a final covariance matrix to test global efficiency and average network strength. Additionally, a range of minimum Pearson's r thresholds between 0.35-0.55 were alternatively applied to edges and binarized to produce a final covariance matrix to test network density. Network construction and measure calculation was performed using the Brain Connectivity Toolbox (Rubinov et al. 2009) in Matlab. Permutation testing in which groups were randomly shuffled 1000 times was used to determine the significance of paired group comparisons for each measure. If significant differences were observed in half or more of the thresholds, they were considered significant. Network measure values for each group are reported as an average across thresholds. These metrics were chosen to index overall brain organization and structural connectivity given extensive evidence of large-scale network alterations in NDDs (Kernbach et al. 2018; Bethlehem et al. 2017).

**Out-of-model Features - Results**

After regressing out the effects of sex, age, and IQ and creating structural covariance networks for each group, average network strength, global efficiency and density were compared using permutation testing.

*Network strength:* On comparison of average network strength across density thresholds, significantly ($p<0.05$) greater average strength was found in Groups 1 (0.67) and 3 (0.66) compared to Groups 2 (0.51). There was also a significant difference between Group 1 and Group 4 (0.59), and between Group 2 versus 4. In contrast, there were no significant differences in average edge strength on comparison of NDD groups (ADHD=0.63, ASD=0.68, OCD=0.76).

*Global network efficiency* did not significantly differ between data-driven (Group 1=0.55, Group 2=0.57, Group 3=0.53, Group 4=0.54) or NDD groups (ASD=0.49, OCD=0.51, ADHD=0.53).

*Network density:* Across Pearson r thresholds, network density was significantly ($p<0.05$) greater in Group 3=0.48 compared to Group 2=0.19; no differences were found between Group 1=0.41 and Group 4=0.32 with any other group. No significant differences in network density were found in NDD groups (ADHD=0.40, ASD=0.54, OCD=0.56) (Figure 3B).

**Cluster Classification**

A random forest machine learning algorithm from the randomForest R package was applied across 100 permutations of randomly resampled participants with a 80/20 train-test split. Participants were proportionally sampled so that in each permutation 80% of each data-driven group was represented. A range of classification models were run to explore which subset of features (when included in the classification model) performed best for prediction of identified data-driven groups (top 10, top 35, all features, or top two features from each of the four data types included in SNF, i.e., cortical thickness, subcortical volume, white matter fractional anisotropy, behavior). Model sensitivity, specificity, overall accuracy, and the Adjusted Rand Index of label overlap are shown in Figure S7. We assessed whether our classification performance was significantly better than chance by comparing it to 500 randomly permuted models with shuffled training labels.

**Table S1. Co-occurring conditions presented by diagnostic and data-driven groups**

| Co-occurring Condition | Diagnostic Groups | | | Data-driven groups | | | |
|---|---|---|---|---|---|---|---|
| | ASD N=30 | ADHD N=49 | OCD N=38 | 1 N=25 | 2 N=43 | 3 N=23 | 4 N=26 |
| Autism Spectrum Disorder | - | 2 | 2 | 1 | 0 | 1 | 2 |
| ADHD (Inattentive, Hyperactivity, Combined) | 11 | - | 8 | 1 | 8 | 5 | 5 |
| OCD | 0 | 1 | - | 0 | 0 | 0 | 1 |
| Anxiety (General, Separation, Social, other) | 9 | 7 | 17 | 9 | 11 | 2 | 11 |
| TIC Disorder | 1 | 3 | 9 | 6 | 1 | 2 | 4 |
| Conduct Disorder or Oppositional Defiant Disorder | 0 | 14 | 2 | 0 | 8 | 3 | 5 |
| IQ<70 | 2 | 0 | 0 | 0 | 1 | 0 | 1 |
| Learning Disorder | 7 | 27 | 2 | 1 | 17 | 9 | 9 |

Comorbidity data was collected using a clinician completed checklist

ADHD=Attention-Deficit/Hyperactivity Disorder

The full scale IQ score was 66 and 62, respectively, for two ASD participants with IQ<70

**Table S2. Sample characteristics presented for the four data-driven groups**

| | 1<br>n=33 | 2<br>n=54 | 3<br>n=41 | 4<br>n=48 | $X_{2df=3}$ | P | post-hoc |
|---|---|---|---|---|---|---|---|
| Sex (F) | 14 (42%) | 15 (21%) | 3 (7%) | 6 (13%) | 17.0 | 0.0007 | 1>2>4>3 |
| Age (years) | 12.3 (2.5) | 11.3 (2.4) | 9.3 (1.8) | 12.1 (2.6) | 33.8 | <0.0001 | 1,2,4>3 |
| FSIQ (WASI & WISC) | 113.7 (15.8) n=17 | 98.0 (16.6) n=47 | 103.1 (13.5) n=31 | 97.9 (18.5) n=41 | 12.4 | 0.01 | 1>2,3,4 |
| Internalizing (CBCL) | 62.7 (8.4) | 62.1 (9.2) | 61.7 (10.9) | 68.8 (7.7) | 16.9 | p<0.0001 | 4>all |
| Externalizing (CBCL) | 50.7 (8.8) | 59.3 (8.3) | 60.7 (11.9) | 62.9 (8.4) | 28.0 | p<0.0001 | 4>2; 2,3,4>1 |
| (TOCS) | 15.5 (16.3) | -15.0 (25.1) | -12.3 (20.9) | 1.0 (23.1) | 39.6 | p<0.0001 | 1>4>2,3, |
| Attention (SWAN) | 0.6 (1.1) | 5.9 (2.3) | 4.7 (2.8) | 5.3 (2.7) | 37.4 | p<0.0001 | 1>2,3,4; 3>2 |
| Hyperactivity (SWAN) | 0.6 (0.9) | 3.1 (2.8) | 4.8 (3.0) | 4.0 (3.0) | 44.7 | p<0.0001 | 1<2,3,4; 2>3 |
| (RBS-R) | 21.0 (15.7) | 18.7 (15.7) | 28.5 (20.5) | 32.1 (23.9) | 13.3 | p<0.0001 | 4>1,2; 3>2 |
| (SCQ) | 5.6 (5.4) | 10.5 (8.1) | 15.6 (9.2) | 18.0 (8.3) | 11.4 | p<0.0001 | 3,4>2>1 |
| (ABAS-II) | 98.1 (16.9) n=30 | 74.5 (17.4) n=52 | 73.2 (16.2) n=37 | 71.8 (16.9) n=48 | 36.4 | p<0.0001 | 1>2,3,4 |

All descriptive statistics are represented as means with standard deviations indicated in brackets except for self-reported sex, where percentage of females is included in brackets. Chi-squared (sex) and Kruskal-Wallis tests (all other measures) were used to test group differences. Post-hoc comparisons were assessed using Dunn's tests. All participants were included in statistics except in the case of IQ and ABAS-II where a subset of the sample was used with available data (as indicated). ASD=Autism Spectrum Disorder, OCD=Obsessive Compulsive Disorder, ADHD=Attention-Deficit/Hyperactivity Disorder, F=female, FSIQ=Full-Scale Intellectual Quotient (estimated using WASI=Wechsler Abbreviated Scale of Intelligence or WISC=Wechsler Intelligence Scale for Children). Internalizing/Externalizing (CBCL)=Child Behaviour Checklist Internalizing or Externalizing t-score. Raw scores for Toronto Obsessive-Compulsive Scale (TOCS), Attention, Hyperactivity (SWAN)=Strengths and Weaknesses of Attention-Deficit/Hyperactivity-symptoms and Normal Behaviours (SWAN) Attention or SWAN Hyperactivity/impulsivity item total scores, RBS-R= Repetitive Behaviours-Revised, SCQ=Social Communication Questionnaire. ABAS-II=The General Adaptive Composite (GAC) Scaled Score for the Adaptive Behaviour Assessment System-II.

**Table S3. All measures included in Similarity Network Fusion and a summary of data-driven group differences (Total = 135)**

| Rank | Top Contributing Features | Normalized Mutual Information | Statistic | pFDR | Eta squared | Post-hoc |
|---|---|---|---|---|---|---|
| **1** | R. pars triangularis CT | 0.182 | 20.3 | 1.06E-09 | 0.315 | 2<1,3,4 |
| **2** | R. insula CT | 0.182 | 24.9 | 4.05E-11 | 0.355 | 2<1,3,4 |
| **3** | SWAN Inattention Score | 0.181 | 16.0 | 3.46E-08 | 0.268 | 1<2,3,4 |
| **4** | R. middle temporal CT | 0.180 | 28.2 | 3.01E-12 | 0.392 | 2<1,3,4 |
| **5** | L. supramarginal CT | 0.172 | 23.7 | 9.67E-11 | 0.343 | 2<1,3,4 |
| **6** | R. inferior temporal CT | 0.169 | 17.5 | 9.36E-09 | 0.285 | 2<1,3,4 |
| **7** | L. middle temporal CT | 0.165 | 29.9 | 1.28E-12 | 0.404 | 2<1,3,4 |
| **8** | R. superior frontal CT | 0.165 | 19.7 | 1.60E-09 | 0.308 | 2<1,3,4 |
| **9** | L. rostral middle frontal CT | 0.162 | 21.7 | 3.68E-10 | 0.321 | 2<1,3,4 |
| **10** | L. insula CT | 0.161 | 21.2 | 5.12E-10 | 0.307 | 2<1,3,4 |
| **11** | L. fusiform CT | 0.159 | 19.3 | 2E-09 | 0.308 | 2<1,3,4 |
| **12** | L. pars opercularis CT | 0.158 | 23.3 | 1.12E-10 | 0.350 | 2<1,3,4 |
| **13** | R. inferior parietal CT | 0.158 | 20.2 | 1.09E-09 | 0.306 | 2<1,3,4 |
| **14** | R. rostral middle frontal CT | 0.157 | 18.0 | 6.33E-09 | 0.292 | 2<1,3,4 |
| **15** | L. inferior parietal CT | 0.157 | 20.4 | 1.0E-09 | 0.304 | 2<1,3,4 |
| **16** | L. inferior temporal CT | 0.153 | 21.8 | 3.68E-10 | 0.335 | 2<1,3,4 |
| **17** | R. precentral CT | 0.149 | 13.2 | 6.66E-07 | 0.228 | 2<1,3,4 |
| **18** | L. lateral orbitofrontal CT | 0.148 | 16.6 | 1.88E-08 | 0.261 | 2<1,3,4 |
| **19** | L. lateral occipital CT | 0.148 | 15.1 | 8.89E-08 | 0.253 | 2<1,3,4 |
| **20** | L. lingual CT | 0.145 | 10.2 | 1.32E-05 | 0.181 | 2<1,3,4 |
| **21** | R. supramarginal CT | 0.145 | 17.2 | 1.24E-08 | 0.272 | 2<1,3,4 |
| **22** | L. pars triangularis CT | 0.137 | 17.5 | 9.36E-09 | 0.281 | 2<1,3,4 |
| **23** | R. fusiform CT | 0.136 | 16.9 | 1.43E-08 | 0.278 | 2<1,3,4 |
| **24** | R. lateral orbitofrontal CT | 0.134 | 13.1 | 6.94E-07 | 0.224 | 2<1,3,4 |
| **25** | L. postcentral CT | 0.134 | 15.2 | 7.8E-08 | 0.259 | 2<1,3,4 |

| | | | | | | |
|---|---|---|---|---|---|---|
| **26** | SWAN Hyperactivity Score | 0.132 | 7.59 | 2.35E-04 | 0.146 | 1<2,3,4 |
| **27** | R. lateral occipital CT | 0.132 | 19.2 | 2.18E-09 | 0.299 | 2<1,3,4 |
| **28** | L. superior frontal CT | 0.131 | 13.9 | 2.91E-07 | 0.239 | 2<1,3,4 |
| **29** | R. medial orbitofrontal CT | 0.127 | 12.9 | 8.3E-07 | 0.217 | 2<1,3,4 |
| **30** | L. pericalcarine CT | 0.126 | 7.49 | 0.000258 | 0.143 | 2<1,3,4 |
| **31** | L. caudal middle frontal CT | 0.123 | 19.6 | 1.69E-09 | 0.310 | 2<1,3,4 |
| **32** | R. caudal middle frontal CT | 0.119 | 18.6 | 3.71E-09 | 0.296 | 2<1,3,4 |
| **33** | R. postcentral CT | 0.118 | 10.4 | 1.01E-05 | 0.194 | 2<1,3,4 |
| **34** | L. pars orbitalis CT | 0.115 | 15.0 | 9.18E-08 | 0.253 | 2<1,3,4 |
| **35** | R. pallidum | 0.115 | 8.02 | 0.000155 | 0.154 | 4<2,3 |
| **36** | L. precentral CT | 0.114 | 17.1 | 1.24E-08 | 0.283 | |
| **37** | L. superior temporal CT | 0.112 | 12.5 | 1.1E-06 | 0.224 | |
| **38** | L. bank of the superior temporal | 0.111 | 18.5 | 3.71E-09 | 0.299 | |
| **39** | L. precuneus CT | 0.111 | 7.81 | 0.0002 | 0.136 | |
| **40** | Total SCQ | 0.107 | 11.4 | 3.8E-06 | 0.192 | |
| **41** | R. putamen volume | 0.106 | 4.21 | 0.0121 | 0.0782 | |
| **42** | L. medial orbitofrontal CT | 0.104 | 7.47 | 0.00026 | 0.126 | |
| **43** | R. superior temporal CT | 0.102 | 11.5 | 3.5E-06 | 0.21 | |
| **44** | R. pars orbitalis CT | 0.101 | 9.8 | 2E-05 | 0.183 | |
| **45** | R. precuneus CT | 0.1 | 10.6 | 8.9E-06 | 0.186 | |
| **46** | R. thalamus volume | 0.099 | 12.6 | 1E-06 | 0.227 | |
| **47** | R. nucleus accumbens volume | 0.0981 | 12.8 | 8.8E-07 | 0.224 | |
| **48** | R. pars opercularis CT | 0.0974 | 12.8 | 8.9E-07 | 0.227 | |
| **49** | R. paracentral CT | 0.0937 | 7.27 | 0.00033 | 0.139 | |
| **50** | R. transverse temporal CT | 0.0927 | 10.9 | 6.3E-06 | 0.198 | |
| **51** | R. bank of the superior temporal | 0.0925 | 7.66 | 0.00022 | 0.145 | |
| **52** | R. lingual CT | 0.092 | 10.9 | 6.3E-06 | 0.18 | |
| **53** | L. thalamus volume | 0.0917 | 12.4 | 1.3E-06 | 0.224 | |
| **54** | Total Obsessive-Compulsive Score | 0.0916 | 8.21 | 0.00013 | 0.154 | |

| | | | | | |
|---|---|---|---|---|---|
| **55** | R. caudate volume | 0.0913 | 7.06 | 0.00041 | 0.136 |
| **56** | R. pericalcarine CT | 0.0898 | 7.64 | 0.00022 | 0.146 |
| **57** | Total Externalizing Score | 0.0879 | 5.6 | 0.00227 | 0.115 |
| **58** | L. cuneus CT | 0.0877 | 7.53 | 0.00025 | 0.136 |
| **59** | L. putamen volume | 0.0873 | 5.3 | 0.00317 | 0.0976 |
| **60** | L. paracentral CT | 0.0862 | 6.39 | 0.0009 | 0.122 |
| **61** | L. superior parietal CT | 0.0842 | 8.76 | 6.5E-05 | 0.158 |
| **62** | L. caudate volume | 0.0837 | 5.69 | 0.00206 | 0.114 |
| **63** | L. nucleus accumbens volume | 0.0827 | 10.8 | 7.2E-06 | 0.19 |
| **64** | L. frontal pole CT | 0.0746 | 9.3 | 3.5E-05 | 0.175 |
| **65** | L. transverse temporal CT | 0.074 | 9.6 | 2.5E-05 | 0.175 |
| **66** | L. rostral anterior cingulate CT | 0.0735 | 1.43 | n.s | |
| **67** | R. isthmus cingulate CT | 0.0726 | 7.67 | 0.00022 | 0.15 |
| **68** | R. superior parietal CT | 0.0717 | 7.74 | 0.00021 | 0.147 |
| **69** | L. hippocampus volume | 0.0688 | 12 | 2.1E-06 | 0.213 |
| **70** | R. frontal pole CT | 0.064 | 6.39 | 0.0009 | 0.129 |
| **71** | L. isthmus cingulate CT | 0.0595 | 4.49 | 0.00853 | 0.0938 |
| **72** | R. para hippocampal CT | 0.0583 | 4.75 | 0.00621 | 0.0981 |
| **73** | R. hippocampus volume | 0.057 | 9.92 | 1.8E-05 | 0.183 |
| **74** | L. posterior cingulate CT | 0.0562 | 3.28 | 0.0358 | 0.0699 |
| **75** | R. cuneus CT | 0.0558 | 3.4 | 0.0316 | 0.0681 |
| **76** | L. para hippocampal CT | 0.0555 | 7.09 | 0.00039 | 0.14 |
| **77** | L. anterior limb of internal FA | 0.0551 | 1.66 | n.s | |
| **78** | L. temporal pole CT | 0.055 | 7.21 | 0.00034 | 0.141 |
| **79** | R. rostral anterior cingulate CT | 0.0548 | 4.18 | 0.0124 | 0.0859 |
| **80** | Total Internalizing Score | 0.0542 | 5.78 | 0.00186 | 0.111 |
| **81** | L. pallidum volume | 0.0528 | 3.49 | 0.0286 | 0.0738 |
| **82** | R. posterior cingulate CT | 0.0513 | 5.57 | 0.00233 | 0.114 |
| **83** | R. retrolenticular internal capsule FA | 0.0476 | 2.68 | n.s | |

| | | | | | |
|---|---|---|---|---|---|
| **84** | R. amygdala volume | 0.0448 | 6.08 | 0.0013 | 0.123 |
| **85** | R. temporal pole CT | 0.0442 | 5.31 | 0.00317 | 0.102 |
| **86** | L. sagittal stratum FA | 0.0434 | 1.13 | n.s | |
| **87** | L. cingulum (hippocampus) FA | 0.0433 | 0.894 | n.s | |
| **88** | R. posterior limb of internal FA | 0.0412 | 3.73 | 0.0215 | 0.0779 |
| **89** | R. cingulum (hippocampus) FA | 0.0406 | 0.826 | n.s | |
| **90** | L. posterior corona radiata FA | 0.0403 | 0.469 | n.s | |
| **91** | L. fornix(cres)/Stria terminalis FA | 0.039 | 1.45 | n.s | |
| **92** | R. cerebral peduncle FA | 0.0385 | 0.85 | n.s | |
| **93** | L. cingulum (cingulate gyrus) FA | 0.0383 | 1.86 | n.s | |
| **94** | R. uncinate fasciculus FA | 0.0357 | 1.32 | n.s | |
| **95** | Fornix (column and body of fornix) | 0.0354 | 2.84 | n.s | |
| **96** | L. caudal anterior cingulate CT | 0.0349 | 0.681 | n.s | |
| **97** | R. cingulum (cingulate gyrus) FA | 0.0347 | 3.2 | 0.0391 | 0.055 |
| **98** | L. amygdala volume | 0.0346 | 4.91 | 0.00517 | 0.0998 |
| **99** | L. external capsule FA | 0.034 | 0.593 | n.s | |
| **100** | R. inferior cerebellar peduncle FA | 0.0336 | 0.0554 | n.s | |
| **101** | L. entorhinal CT | 0.0328 | 3.73 | 0.0215 | 0.0789 |
| **102** | R. sagittal stratum FA | 0.0326 | 1.06 | n.s | |
| **103** | L. cerebral peduncle FA | 0.0317 | 0.196 | n.s | |
| **104** | R. caudal anterior cingulate FA | 0.0304 | 1.2 | n.s | |
| **105** | L. corticospinal tract FA | 0.029 | 0.715 | n.s | |
| **106** | R. entorhinal CT | 0.0285 | 3.54 | 0.0269 | 0.0708 |
| **107** | R. anteriorlimbinternalcapsuleFA | 0.0283 | 0.339 | n.s | |
| **108** | L. uncinate fasciculus FA | 0.0279 | 1.98 | n.s | |
| **109** | L. superior longitudinal fasciculus | 0.0276 | 0.331 | n.s | |
| **110** | L. anterior corona radiata FA | 0.0273 | 0.315 | n.s | |
| **111** | Total Repetitive Behaviours Score | 0.0272 | 3.32 | 0.0345 | 0.0681 |
| **112** | R. medial lemniscus FA | 0.027 | 1.22 | n.s | |

| | | | | |
|---|---|---|---|---|
| **113** | R. superior longitudinal fasciculus | 0.0267 | 0.415 | n.s |
| **114** | L. inferior cerebellar peduncle FA | 0.0263 | 0.209 | n.s |
| **115** | R. superior cerebellar peduncle FA | 0.0255 | 1.29 | n.s |
| **116** | L. superior corona radiata FA | 0.0253 | 0.953 | n.s |
| **117** | L. posterior limb of internal capsule | 0.0242 | 2.76 | n.s |
| **118** | L. retrolenticularinternalcapsule FA | 0.0242 | 1.86 | n.s |
| **119** | R. posterior corona radiata FA | 0.024 | 0.484 | n.s |
| **120** | Genu of corpus callosum FA | 0.0239 | 0.606 | n.s |
| **121** | R. posterior thalamic radiation FA | 0.0238 | 1.06 | n.s |
| **122** | R. external capsule FA | 0.0234 | 0.11 | n.s |
| **123** | R. inferior fronto-occipital fasciculus | 0.0231 | 0.79 | n.s |
| **124** | R. superior corona radiata FA | 0.0203 | 0.695 | n.s |
| **125** | Body of corpus callosum FA | 0.0203 | 0.523 | n.s |
| **126** | L. medial lemniscus FA | 0.02 | 0.184 | n.s |
| **127** | L. inferior fronto-occipital fasciculus | 0.0175 | 0.244 | n.s |
| **128** | L. superior fronto-occipital | 0.0169 | 0.91 | n.s |
| **129** | R. anterior corona radiata FA | 0.0162 | 0.102 | n.s |
| **130** | Splenium of corpus callosum FA | 0.0153 | 1.4 | n.s |
| **131** | R. superior fronto-occipital | 0.0135 | 0.87 | n.s |
| **132** | R. Fornix(cres)/Stria terminalis FA | 0.0124 | 0.2 | n.s |
| **133** | R. corticospinal tract FA | 0.0117 | 1.22 | n.s |
| **134** | L. superior cerebellar peduncle FA | 0.00909 | 1.54 | n.s |
| **135** | L. posterior thalamic radiation FA | 0.00797 | 0.0523 | n.s |

R.=right, L.=left, CT=cortical thickness, FA=fractional anisotropy, Repetitive Behaviors score=Repetitive Behaviors Scale-Revised, total social communication=total Social Communication Questionnaire score, SWAN inattention=Strengths and Weaknesses of Attention-Deficit/Hyperactivity-symptoms and Normal Behaviors (SWAN) inattention items score, SWAN= Strengths and Weaknesses of Attention-Deficit/Hyperactivity-symptoms and Normal Behaviours (SWAN) hyperactivity items score, Total obsessive compulsive score = Toronto Obsessive-Compulsive Scale total score, total internalizing= Child Behavior Checklist Internalizing, total externalizing= Child Behavior Checklist Externalizing. Group differences were determined using ANOVAs while including age, sex and IQ as covariates. Post-hoc comparisons were assessed using Tukey tests.

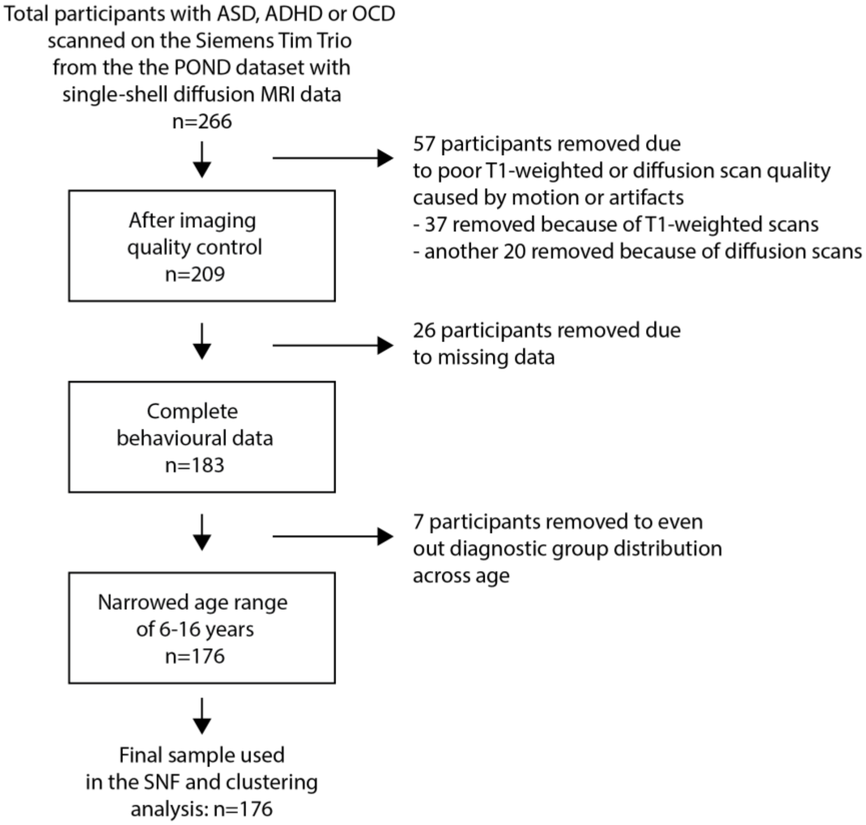


Figure S1. Flowchart of total participants and how many were removed at each level of quality control for the current analysis

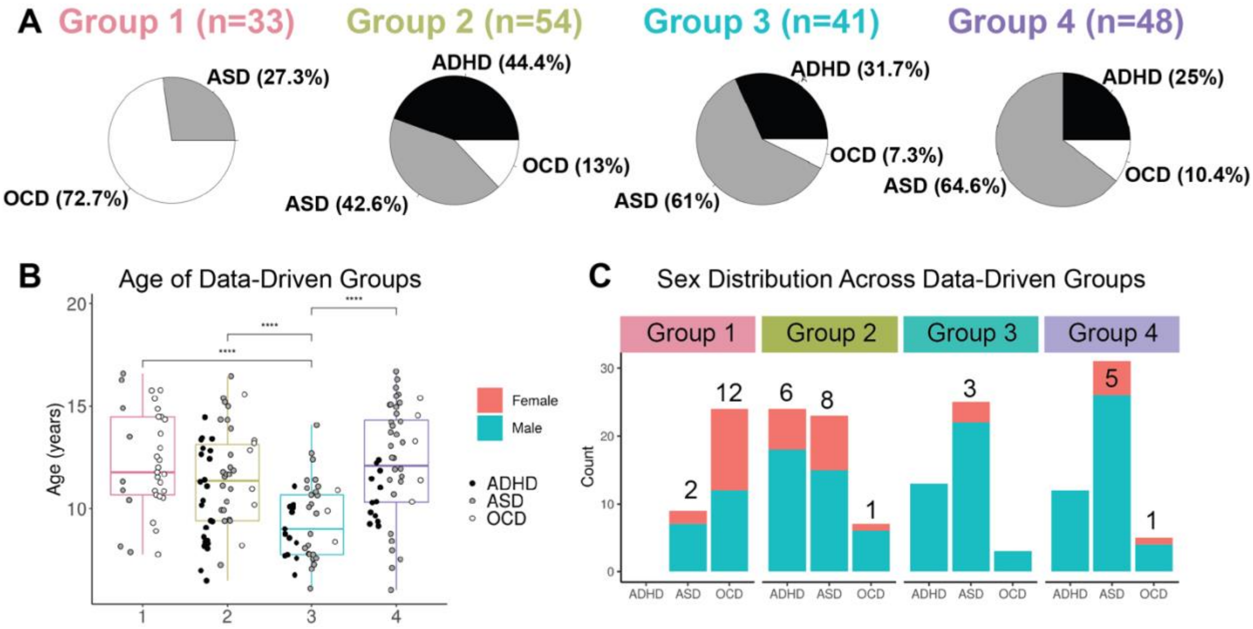


Figure S2. (A) Breakdown of clinical diagnoses within each data-driven group. (B) Age differences between data-driven groups. The boxplot shows the first and third quartile with extensions representing 1.5 * the inter-quartile range. (C) Distribution of males and females in the data-driven groups by neurodevelopmental disorder.

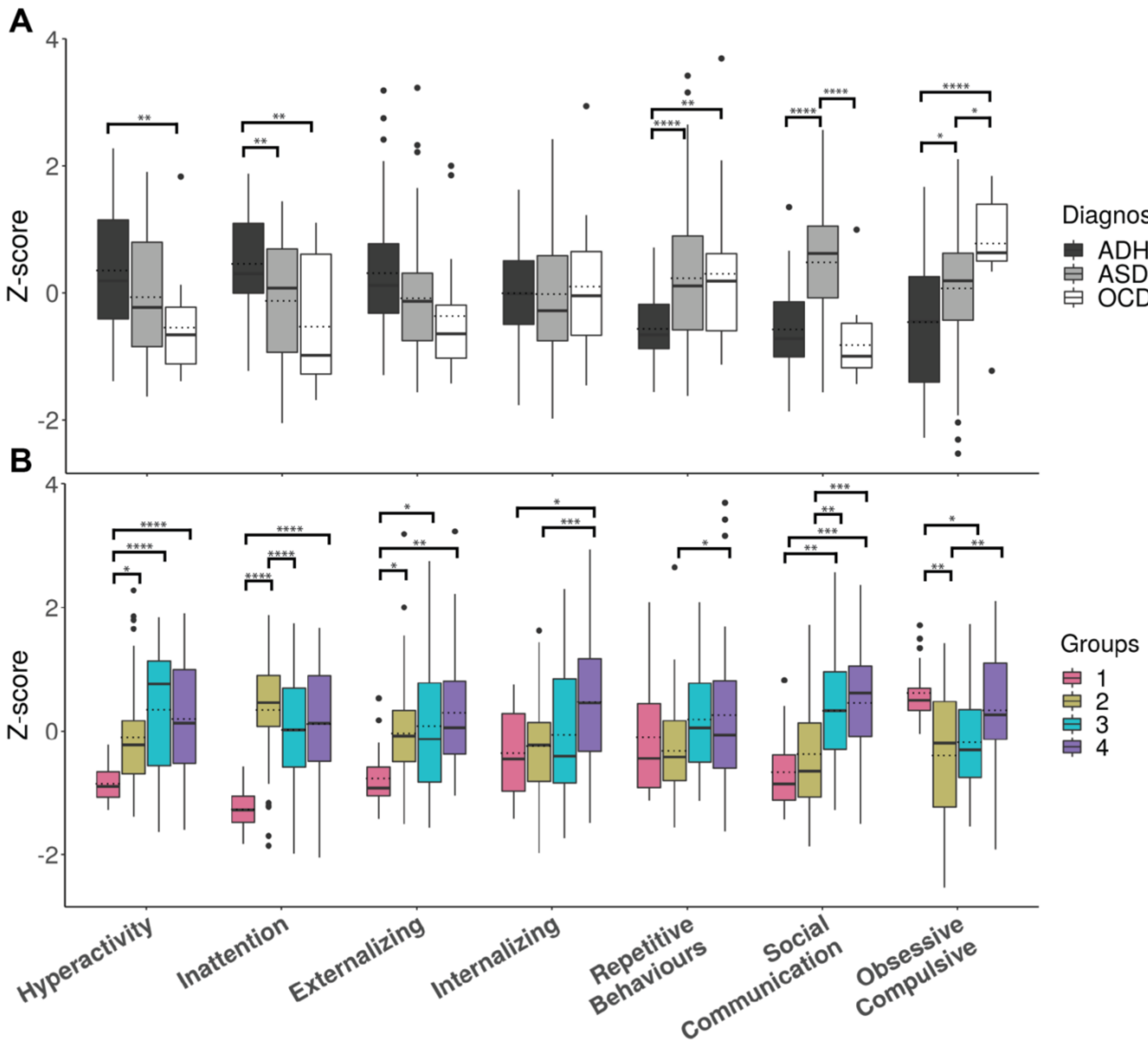


Figure S3. Behavioral scores for both (A) neurodevelopmental disorders (NDD) groups (i.e., ASD=Autism Spectrum Disorder, OCD=Obsessive Compulsive Disorder, ADHD=Attention-Deficit/Hyperactivity Disorder), and (B) data-driven groups after the effects of age, sex and IQ have been regressed out. Hyperactivity = Strengths and Weaknesses of Attention-Deficit/Hyperactivity-symptoms and Normal Behaviors (SWAN) hyperactivity/impulsivity symptoms, Inattention= SWAN inattention symptoms, Externalizing or Internalizing= Child Behavior Checklist (CBCL) Internalizing or Externalizing broad-band raw score. Repetitive Behaviors= Repetitive Behaviors-Revised total score, Social Communication=Social Communication Questionnaire total score, Raw scores for Toronto Obsessive-Compulsive Scale (TOCS). Boxplots show the first and third quartile with extensions representing 1.5 * the inter-quartile range. Dashed lines indicate group means and solid lines indicate medians. Error bars show standard errors. Stars indicate significance on follow-up Tukey tests (****<0.00005, ***<0.0005, **<0.005, *<0.05).

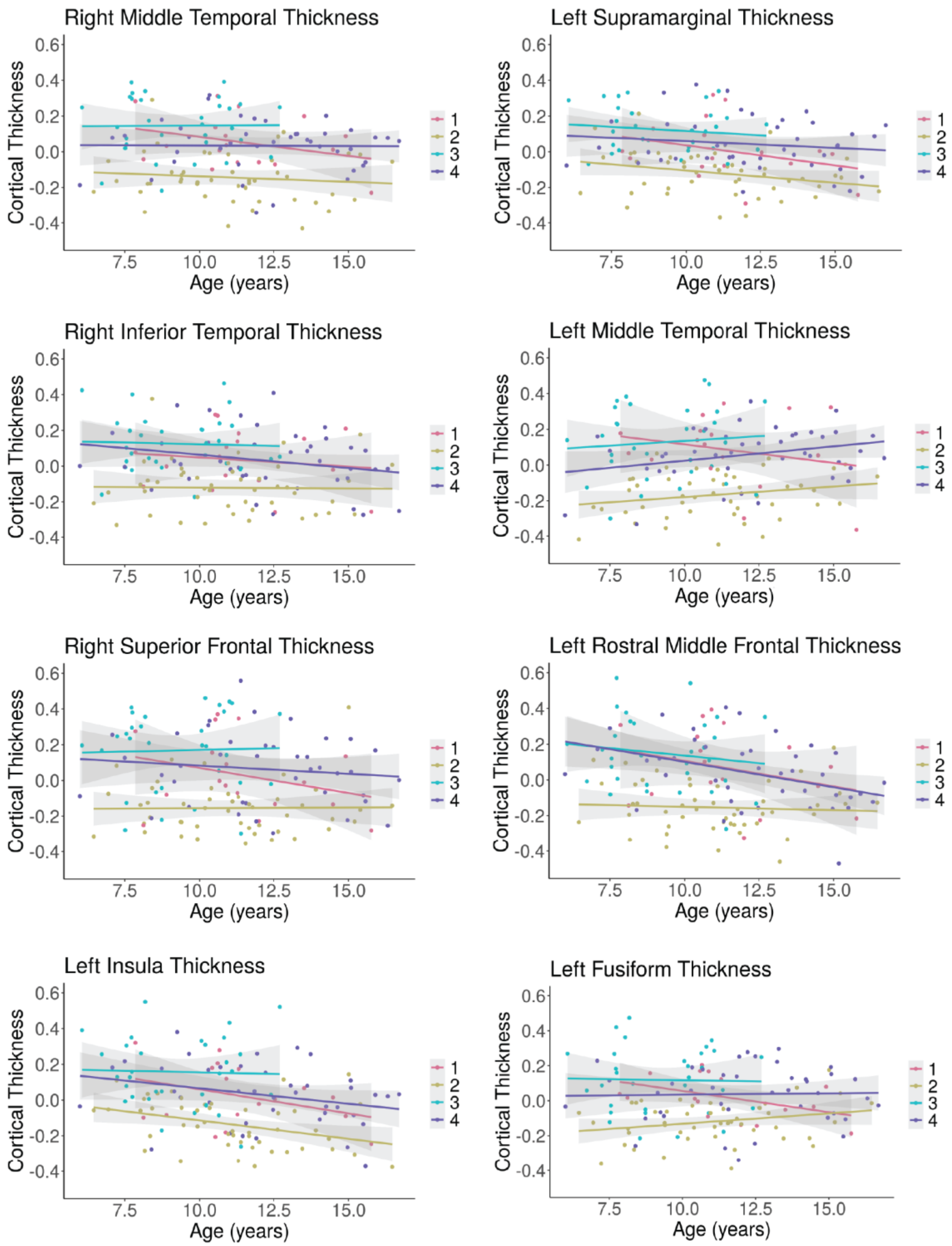

Figure S4. Top cortical thickness regions contributing to participant similarity presented across age. Graphs show values after the effects of sex and IQ have been regressed out. Shaded areas represent 95% confidence intervals.

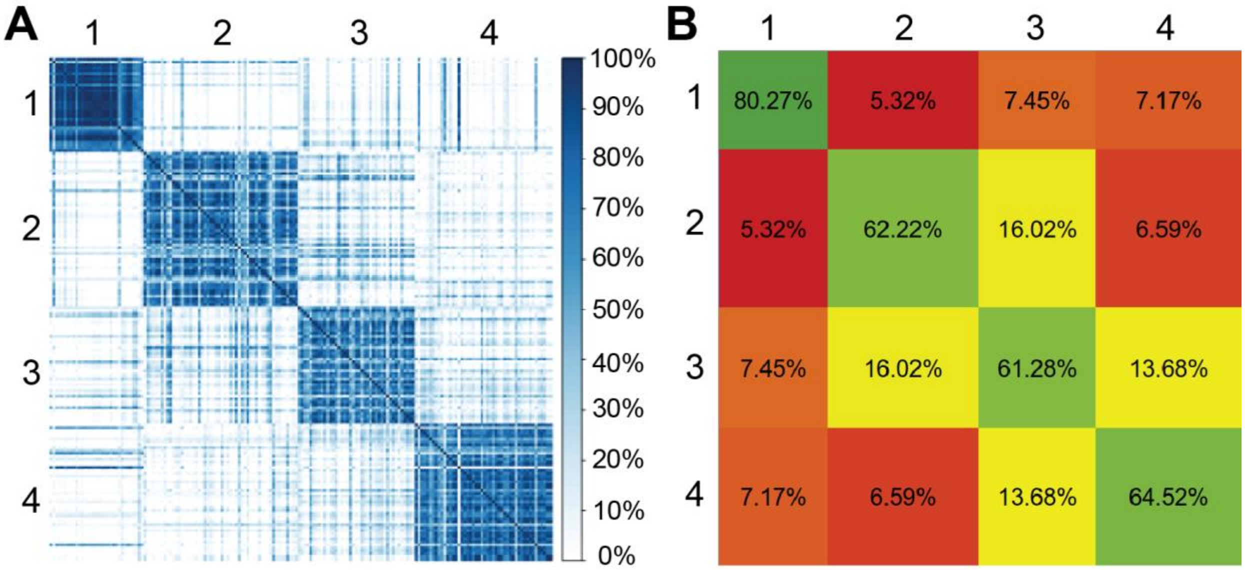


Figure S5. Two representations of the participant agreement matrix (how often each participant clustered with each other participant across resampling). (A) Darker blue indicates a higher percentage of instances that each pair of participants clustered together across resampling 80% of participants 1000 times. (B) Average percentage of how often each participant clusters with each other participant within each data-driven group (shown on the diagonal in green) or with other groups across resampling. Groups 1-4 are presented across from left to right and top to bottom.

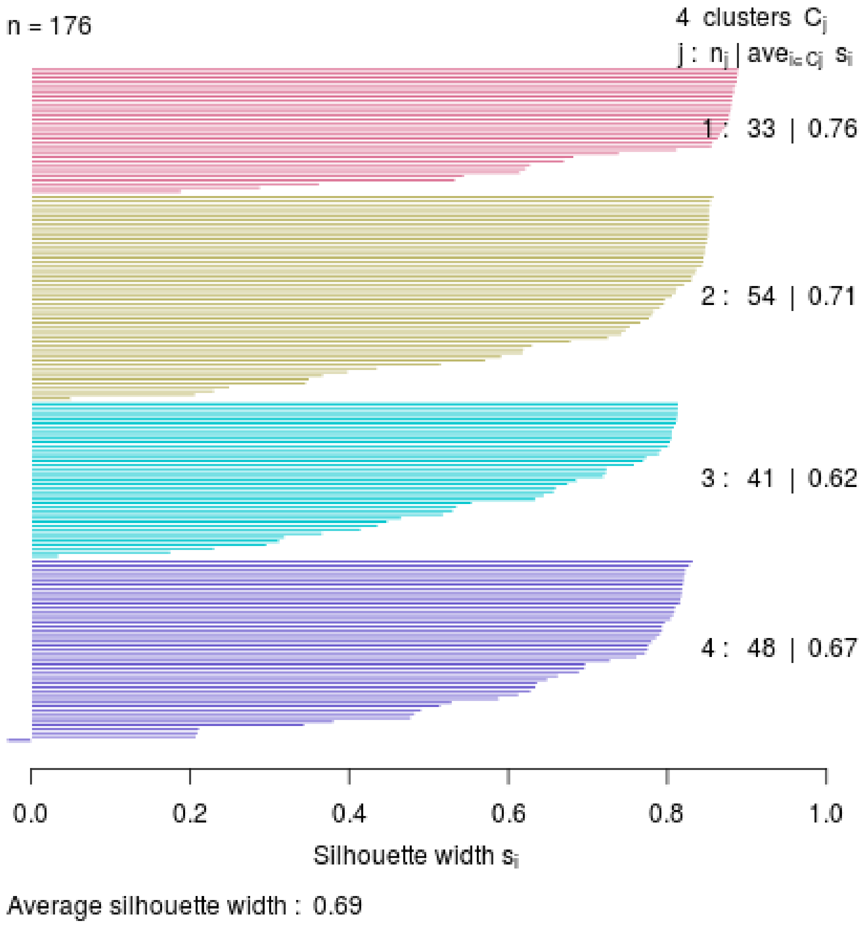


Figure S6. Silhouette plot illustrating the similarities between each participant and the other participants within their group as compared to similarities among participants in other groups. Group number: Group n (i.e., number of participants in each group) | average silhouette width for each group, are indicated on the right side of the plot. The R package cluster v2.0.7 was used to create this figure.

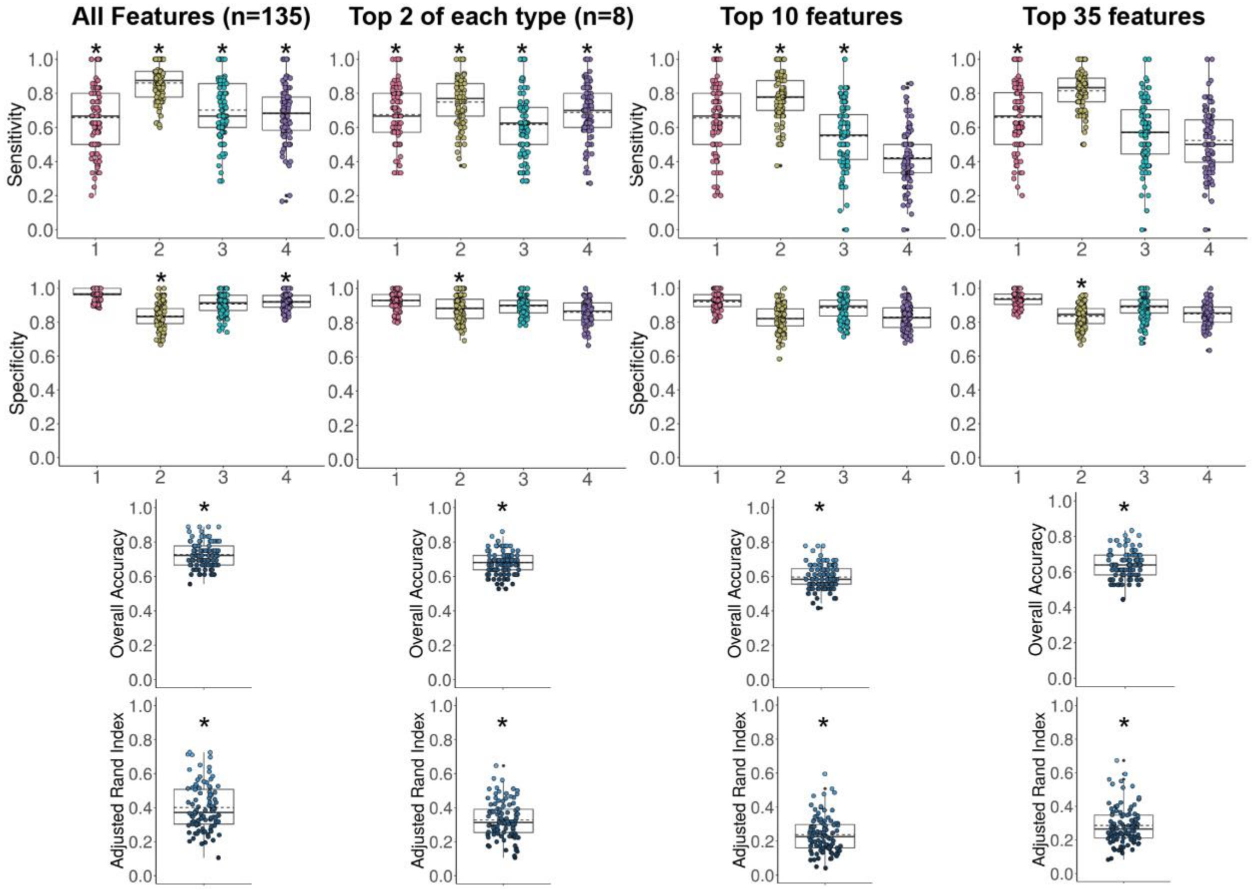


Figure S7. Model sensitivity, specificity, overall model accuracy, and adjusted rand index values between predicted and true data-driven group labels are presented to demonstrate random forest classifier performance across 100 models. Different classifier models tested included: all features included in the SNF model, the top two contributing features within each data type, top 10, and top 35 contributing features. A star indicates that performance is significantly better than chance ($p<0.05$) as determined using randomly permuted models. Boxplots show the first and third quartile with extensions representing 1.5 * the inter-quartile range. The dotted lines represent the mean and the solid lines represent the median.

**References**


1. Andersson, Jesper L. R., Mark S. Graham, Enikő Zsoldos, and Stamatios N. Sotiropoulos. 2016. “Incorporating Outlier Detection and Replacement into a Non-Parametric Framework for Movement and Distortion Correction of Diffusion MR Images.” *NeuroImage* 141 (November): 556–72.
2. Andersson, Jesper L. R., and Stamatios N. Sotiropoulos. 2016. “An Integrated Approach to Correction for off-Resonance Effects and Subject Movement in Diffusion MR Imaging.” *NeuroImage* 125 (January): 1063–78.
3. Bethlehem, R. A. I., R. Romero-Garcia, E. Mak, E. T. Bullmore, S. Baron-Cohen, and Richard A. I. Bethlehem. 2017. “Structural Covariance Networks in Children with Autism or ADHD.” *Cerebral Cortex* 27: 4267–76.
4. Dietrich, Olaf, José G. Raya, Scott B. Reeder, Maximilian F. Reiser, and Stefan O. Schoenberg. 2007. “Measurement of Signal-to-Noise Ratios in MR Images: Influence of Multichannel Coils, Parallel Imaging, and Reconstruction Filters.” *Journal of Magnetic Resonance Imaging: An Official Journal of the International Society for Magnetic Resonance in Medicine* 26 (2): 375–85.
5. Esteban, Oscar, Daniel Birman, Marie Schaer, Oluwasanmi O. Koyejo, Russell A. Poldrack, and Krzysztof J. Gorgolewski. 2017. “MRIQC: Advancing the Automatic Prediction of Image Quality in MRI from Unseen Sites.” *PloS One* 12 (9): e0184661.
6. Esteban, Oscar, Christopher J. Markiewicz, Ross W. Blair, Craig A. Moodie, A. Ilkay Isik, Asier Erramuzpe, James D. Kent, et al. 2019. “fMRIPrep: A Robust Preprocessing Pipeline for Functional MRI.” *Nature Methods* 16 (1): 111–16.
7. Ganzetti, Marco, Nicole Wenderoth, and Dante Mantini. 2016. “Intensity Inhomogeneity Correction of Structural MR Images: A Data-Driven Approach to Define Input Algorithm Parameters.” *Frontiers in Neuroinformatics* 10 (March): 10.
8. Hubert, Lawrence, and Phipps Arabie. 1985. “Comparing Partitions” 218: 193–218.
9. Kernbach, Julius M., Theodore D. Satterthwaite, Danielle S. Bassett, Jonathan Smallwood, Daniel Margulies, Sarah Krall, Philip Shaw, et al. 2018. “Shared Endo-Phenotypes of Default Mode Dsfunction in Attention Deficit/hyperactivity Disorder and Autism Spectrum Disorder.” *Translational Psychiatry* 8 (1). https://doi.org/10.1038/s41398-018-0179-6.

10. Magnotta, Vincent A., Lee Friedman, and FIRST BIRN. 2006. “Measurement of Signal-to-Noise and Contrast-to-Noise in the fBIRN Multicenter Imaging Study.” *Journal of Digital Imaging* 19 (2): 140–47.
11. Pardoe, Heath R., Rebecca Kucharsky Hiess, and Ruben Kuzniecky. 2016. “Motion and Morphometry in Clinical and Nonclinical Populations.” *NeuroImage* 135 (July): 177–85.
12. Rubinov, M., R. Kötter, P. Hagmann, and O. Sporns. 2009. “Brain Connectivity Toolbox: A Collection of Complex Network Measurements and Brain Connectivity Datasets.” *NeuroImage*. https://doi.org/10.1016/s1053-8119(09)71822-1.
13. Wang, Bo, Aziz M. Mezlini, Feyyaz Demir, Marc Fiume, Zhuowen Tu, Michael Brudno, Benjamin Haibe-Kains, and Anna Goldenberg. 2014. “Similarity Network Fusion for Aggregating Data Types on a Genomic Scale.” *Nature Methods* 11 (January): 333.
14. Zalesky, Andrew, Alex Fornito, and Edward T. Bullmore. 2010. “Network-Based Statistic: Identifying Differences in Brain Networks.” *NeuroImage* 53 (4): 1197–1207.
